# Primary myeloid cell proteomics and transcriptomics: importance of ß tubulin isotypes for osteoclast function

**DOI:** 10.1101/800516

**Authors:** David Guérit, Pauline Marie, Anne Morel, Justine Maurin, Christel Verollet, Brigitte Raynaud-Messina, Serge Urbach, Anne Blangy

## Abstract

Among hematopoietic cells. osteoclasts (Oc) and immature dendritic cells (Dc) are closely related myeloid cells with distinct functions; Oc participate skeleton maintenance while Dc sample the environment for foreign antigens. Such specificities rely on profound modifications of gene and protein expression during Oc and Dc differentiation. We provide global proteomic and transcriptomic analyses of primary mouse Oc and Dc. based on original SILAC and RNAseq data. We established specific signatures for Oc and Dc including genes and proteins of unknown functions. In particular. we showed that Oc and Dc have the same α and β tubulin isotypes repertoire but that Oc express much more β tubulin isotype Tubb6. In both mouse and human Oc. we demonstrate that elevated expression of Tubb6 in Oc is necessary for correct podosomes organization and thus for the structure of the sealing zone. which sustains the bone resorption apparatus. Hence. lowering Tubb6 expression hindered Oc resorption activity. Overall. we highlight here potential new regulators of Oc and Dc biology and illustrate the functional importance of the tubulin isotype repertoire in the biology of differentiated cells.

**Summary statement:** This study provides original proteomic and transcriptomic data of primary myeloid cells. The analysis led to signatures for osteoclasts and for immature dendritic cells including potential new regulators of their specific biology. RNA interference showed in particular that ß tubulin isotype Tubb6 participates in osteoclast podosome patterning. sealing zone structure and in the resorption activity.

## Introduction

Osteoclasts (Oc) and monocyte-derived immature dendritic cells (Dc) are closely related cell types belonging to the monocyte lineage of myeloid hematopoietic cells. Although closely related and capable of trans-differentiation (Madel et al.. 2019). Oc and Dc fulfill very distinct biological functions. On one hand. Oc reside in the bone tissue and are specialized for bone resorption; they ensure the maintenance of bone health in coordination with osteoblasts. the bone forming cells. Oc function is also crucial for hematopoietic stem cell mobilization and they are closely linked with the immune system (Cappariello et al.. 2014). On the other hand. immature dendritic cells are sentinels of the immune system that reside in peripheral tissues where they constantly sample the surrounding environment for pathogens. such as viruses or bacteria; upon non-self body encountering. Dc mature or differentiate into antigen presenting cells to fulfill their immune-initiating function (Tiberio et al.. 2018). Myeloid cell differentiation is associated with the expression of distinctive proteins related to their specific functions. such as the Cathepsin K and the metalloprotease MMP9 in Oc for bone collagen fiber degradation (Cappariello et al.. 2014) or the chemokine CCL17 in Dc for their migration and T cell chemo-attraction (Real et al.. 2004). In particular. Oc have a very specific capacity to organize their actin cytoskeleton into a belt of podosomes. which is the backbone of the bone resorption apparatus (Georgess et al.. 2014a; Touaitahuata et al.. 2014). whereas Dc display clusters of podosomes that participate in antigen sampling (Baranov et al.. 2014).

Unraveling the fine molecular mechanisms controlling the specific functions of Oc and Dc is key to understand the homeostasis and pathological dysfunctions of the bone and immune system homeostasis. and then to design targeted genetic and pharmacological therapeutic strategies. In fact. improving knowledge about Dc biology and pathology paved the way for the manipulation of Dc to enhance immune responses. in particular in the context of cancer immunotherapy (Sabado et al.. 2017; Wculek et al.. 2019). The exacerbated activity of Oc causes osteoporosis. a serious public health problem leading to significant morbidity and mortality; this abnormal increase in Oc activity is associated with a number of pathologies ranging from age-related sexual hormone decay to chronic inflammation and cancer (Khosla and Hofbauer. 2017). Better understanding of Oc biology allowed to envision more focused therapeutic solutions against osteoporosis. as demonstrated by the recent approval of Denosumab. an antibody against the Receptor Activator of NF-κB Ligand (RANKL). the key cytokine for Oc differentiation (Compston et al.. 2019) and the promising targets Cathepsin K (Drake et al.. 2017) or Dock5 (Vives et al.. 2015).

Differential proteomics and transcriptomics proved powerful tools to identify molecular pathways involved in specific cell functions. The comparison between Oc and Dc transcripts indeed revealed new regulators of bone resorption by Oc (Gallois et al.. 2010; Georgess et al.. 2014b) and combined Oc transcriptomics and proteomics shed light to the changes in cellular functions associated with Oc differentiation (An et al.. 2014). Nevertheless. the very few studies available in myeloid cells are difficult to compare due to different cellular systems and experimental set up. Furthermore. the only proteomics data available for Oc were obtained with Oc derived from the RAW264.7 cell line. which proved to be very different from primary Oc. in particular regarding cytoskeleton regulation (Ng et al.. 2018) that is key for bone resorption (Georgess et al.. 2014a; Touaitahuata et al.. 2014) and a promising therapeutic target against osteoporosis (Vives et al.. 2015).

To overcome the limitation of currently available data. we provide here comparative quantitative proteomics and transcriptomics data of primary Oc. Dc and bone marrow macrophages (Mo). differentiated *ex vivo* from the same bone marrow cells. By minimalizing the culture condition differences to the cytokine cocktails used to differentiate each myeloid cell type. we could establish transcriptomic and proteomic signatures for Oc and Dc. We then exploited the data to uncover potential new regulators of Oc biology and highlighted the ß-tubulin isotype Tubb6 as key for Oc cytoskeleton organization and bone resorption capacity.

## Results

### Global proteome and transcriptome of primary myeloid cells

Osteoclasts (Oc) and immature dendritic cells (Dc) are myeloid cells that perform very distinct biological functions. In order to get new insight into the molecular mechanisms underlying their specific activities. we sought to compare the global proteome and transcriptome of primary Oc and Dc. which are post mitotic myeloid cells. using bone marrow macrophages (Mo). which retain the ability to divide in our experimental conditions. as reference monocytic myeloid cell type. For this and to minimize irrelevant variations due to culture conditions. we set up conditions to derive the three myeloid cell types from the same mouse bone marrow cells with the only variation in culture medium composition being restricted to cytokines: GM-CSF for Dc. M-CSF for Mo and M-CSF + RANKL for Oc (Fig. 1A-B). In each experiment. we monitored Dc and Oc differentiation during and at the end of differentiation by controlling cell morphology and the cytoskeleton using phase-contrast and fluorescence microscopy (Fig. S1A-B).

**FIG. 1.**
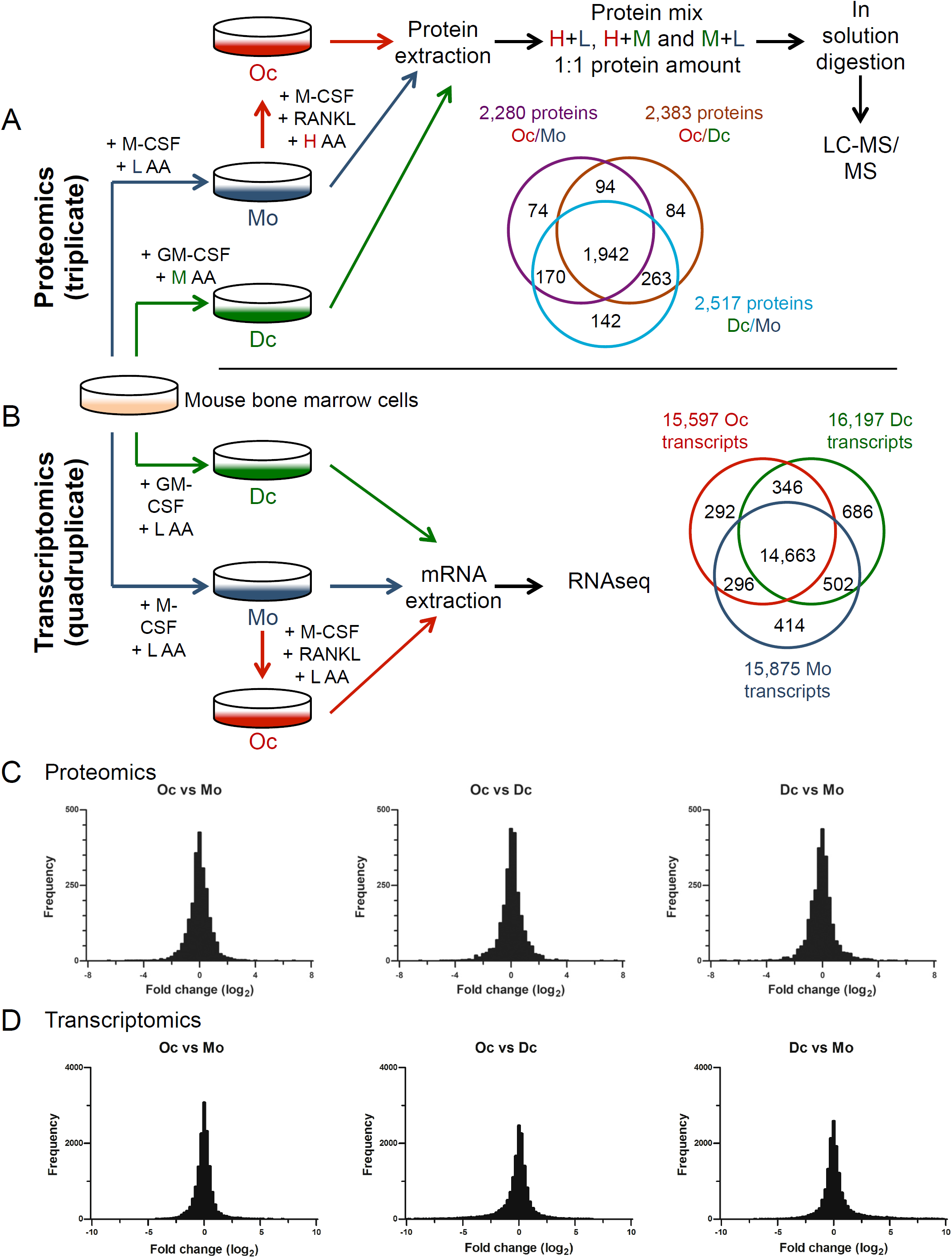
Proteomics and transcriptomics of primary myeloid cells. **(A-B)** Experimental procedure scheme for SILAC (A) and RNAseq (B) in primary myeloid cells: bone marrow cells were cultured in presence of M-CSF or GM-CSF to obtain Mo and Dc respectively. Oc were obtained by adding RANKL to the Mo cultures. H. M and L isotopic-labeled amino acids were used for Oc. Dc and Mo respectively for SILAC whereas unlabeled amino acid were used for RNAseq. Venn diagrams indicate the distribution of proteins in SILAC for Oc vs Mo. Oc vs Dc and Dc vs Mo comparisons and the distribution of Oc. Dc and Mo transcripts in RNAseq. **(C-D**) Histograms show the distribution of log_2_ fold changes in proteome (C) and transcriptome (D) for the indicated comparisons (bin size = 0.25).

To quantify proteins. we performed SILAC with isotope-labeled Arg and Lys: heavy-isotopic labeling (H) for Oc. intermediate-isotopic labeling (M) for Dc and no isotopic labeling (L) for Mo (Fig. 1A). incorporating isotope-labeled aminoacids at the time of cytokines addition in order to label only proteins neosynthetized during Dc and Oc differentiation. Protein levels were compared between Oc versus (vs) Mo (H+L). Dc vs Mo (M+L) and Oc vs Dc (H+M). To quantify transcripts. we performed RNAseq in identical culture conditions as SILAC except for labeled amino acids (Fig. 1B). Principal component analysis showed that SILAC and RNAseq biological replicates were grouped whereas the 3 SILAC clusters (H+L. M+L and H+M) and the 3 RNAseq clusters (Oc. Dc and Mo) were clearly separate (Fig. S2A-B). as confirmed by multi scatter plotting (Fig. S3-4). We further used these global SILAC and RNAseq data to identify the proteomic and transcriptomic specificities of primary Oc and Dc.

SILAC identified a total of 2.820 protein groups considering one peptide for adequate protein identification. among which 2.769 corresponded unambiguously to a single UniProt identifier and to a single protein (Table S1). Among these. 1.942 were present in all three myeloid cell comparisons. the others only in one or two (Fig. 1A). The normalized log_2_ protein abundance ratios were in the same range in all comparisons. between around −7 and +7 (Fig. 1C and Table S1). Considering only genes annotated as “protein coding” in the Ensembl database. the RNAseq identified 15.597 mouse Ensembl genes in Oc. 16.197 in Dc and 15.875 in Mo. representing collectively 17.199 different genes. with most of the transcripts (14.663) present in the three myeloid cell types (Fig. 1B and Table S2). Differential expression analysis of the 4 replicate experiments picked 4.781 significant transcript differences in Oc vs Mo. 6.903 in Oc vs Dc and 6.766 in Dc vs Mo (padj < 0.05 in Table S2). The log_2_ fold changes were in the same range for in all comparisons. between roughly −10 and +10 (Fig. 1D and Table S2). For gene expression profiling. we excluded the 7.624 genes of similar expression levels in Oc. Dc and Mo (italic in Table S2) and the 540 genes with a mean expression level below 10 Fragments Per Kilobase Million (FPKM) in the three cell types (italic in Table S2). Hierarchical clustering analysis of the remaining 9.035 genes revealed 5 gene clusters. in particular cluster E corresponding to elevated expression in Oc and clusters A and C to elevated expression in Dc (Fig. 2).

**FIG. 2.**
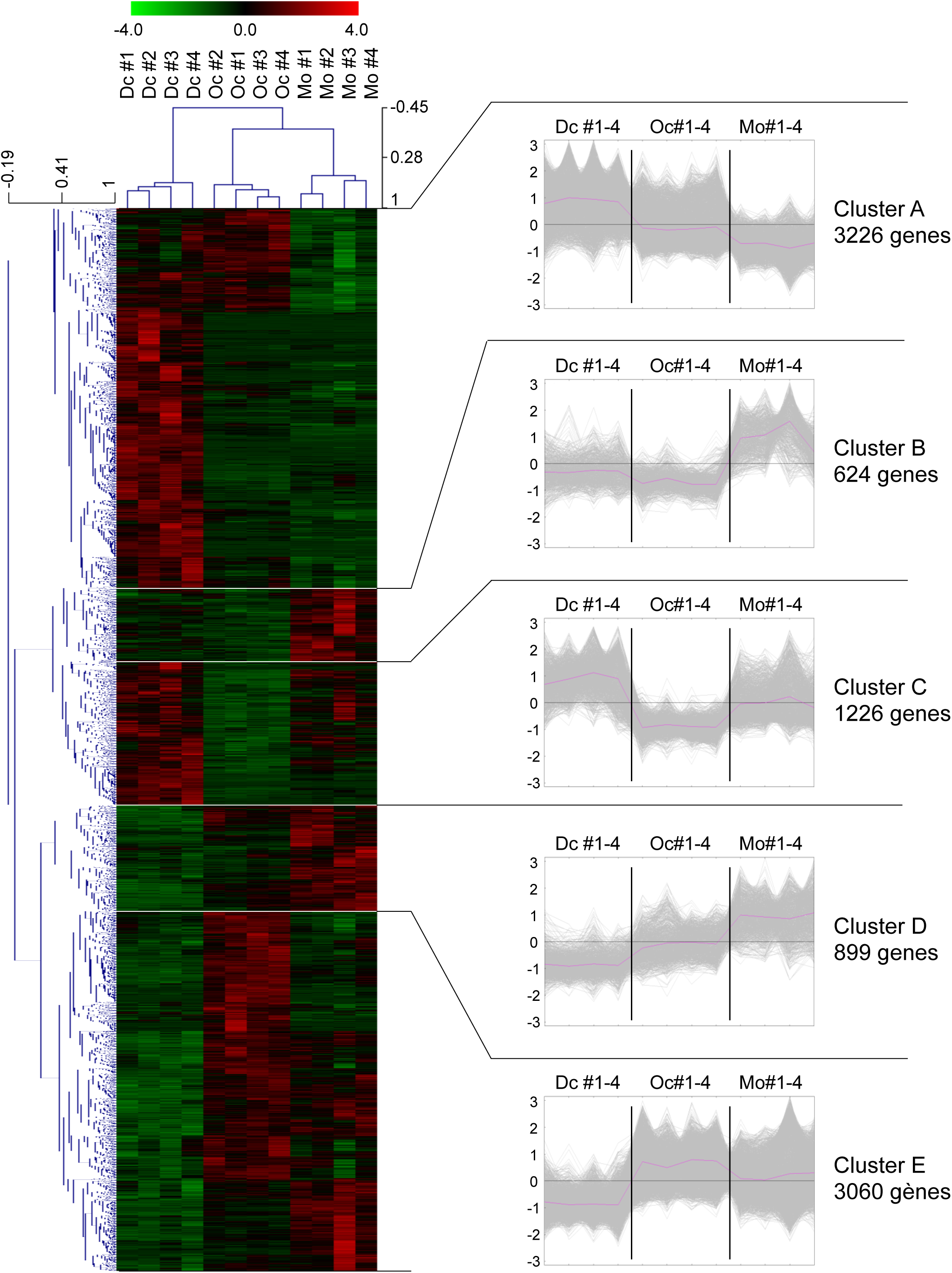
Hierarchical clustering of genes differentially expressed in primary myeloid cells. Hierarchical clustering of the 12 RNAseq samples (columns) with gene expression heat map of the 9.035 genes (lines) differentially expressed between Oc. Mo and Dc (according to the data in Table S2) with the corresponding genes expression profile clusters on the right.

### Analysis of differential protein and gene expression between myeloid cells

We then identified proteins and genes characteristic of Oc and of Dc. For proteins. expression levels were considered different when the normalized log_2_ abundance ratio was ≥ 0.75 or ≤-0.75. Thereby. we established an Oc protein signature of 144 proteins. present at lower levels in both Dc and Mo (Fig. 3A and Table S3). Similarly. the immature Dc signature comprised 181 proteins (Fig. 3A and Table S4). Conversely. 80 proteins were less abundant in Oc (Fig. 3A and Table S5) and 147 proteins less abundant in Dc (Fig. 3A and Table S6). A similar analysis for the transcripts analyzed by hierarchical clustering led to Oc and Dc transcriptional signatures of 1.207 and 2.419 genes respectively (Fig. 3B and Table S2). Finally. 554 genes were less expressed in Oc (Fig. 3B and Table S2) and 1.587 genes less expressed in Dc (Fig. 3B and Table S2).

**FIG. 3.**
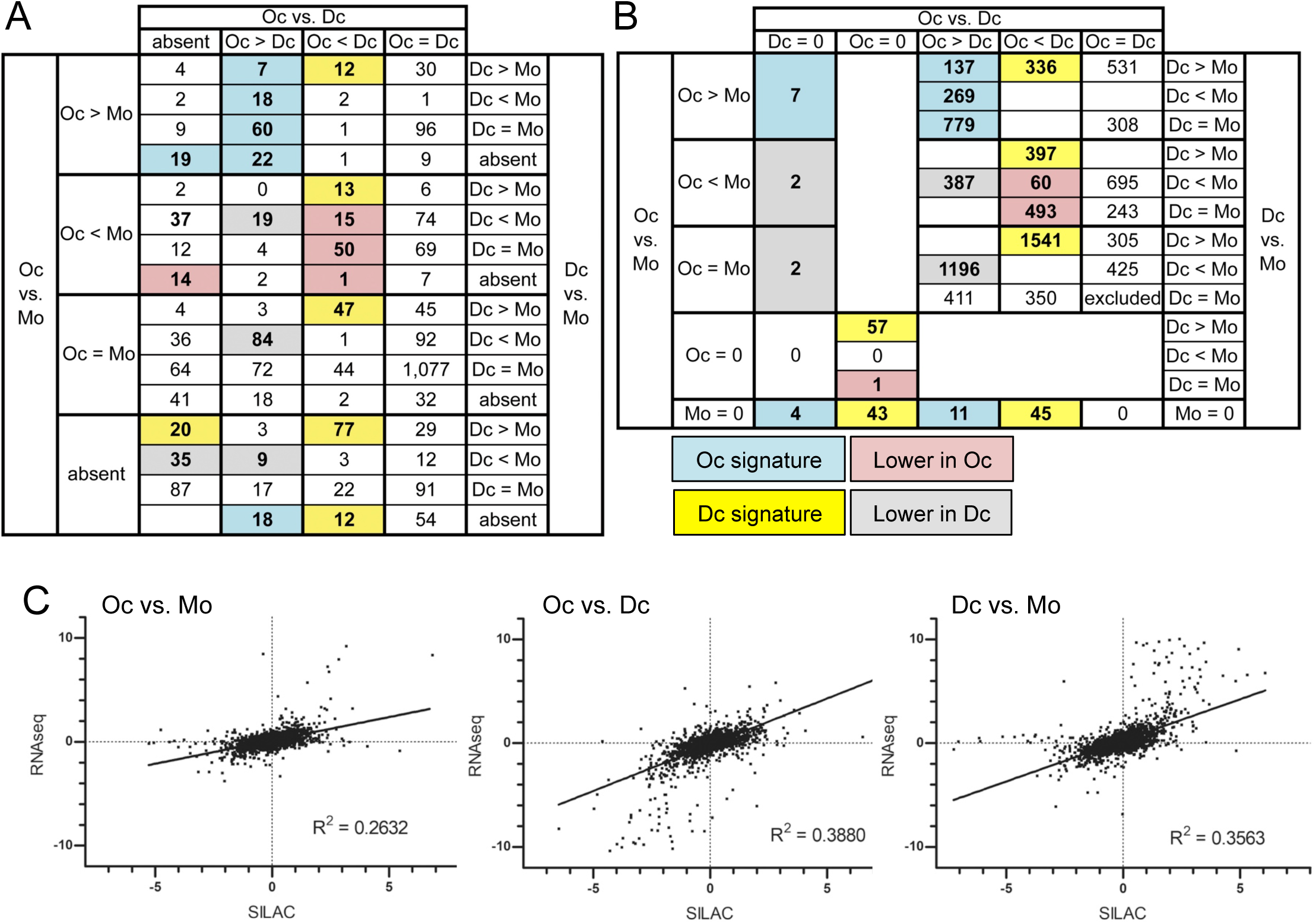
Global transcriptomic and proteomic data analysis to establish the Oc and Dc signatures. **(A-B)** Comparative expression profiles of proteins (A) and genes (B) in Oc. Dc and Mo. Colored boxes are the differentially expressed proteins and genes counts. the details of the proteins are in Tables S3 and S5 for Oc and Tables S4 and S6 for Dc. the details of the genes are in Table S2 with the same color code. In blue: more expressed in Oc (Oc signatures): 7+18+60+22+19+18=144 proteins and 137+269+779+7+11+4=1.207 genes. in yellow: more expressed in Dc (Dc signatures): 12+13+47+77+20+12=181 proteins and 336+397+1541+57+45+43=2.419 genes; in red: genes and proteins less expressed in Oc and in grey: genes and proteins less expressed in Dc. **(C)** Correlation between proteomic and transcriptomic studies: for each mRNA-protein pair and each cell type comparison. graphs show the log_2_ abundance ratio from SILAC in the x-axis and the corresponding log_2_ fold change value from RNAseq in the y-axis. R2: correlation coefficients.

R^2^ correlation coefficients between proteomics and transcriptomics were low around 0.3 (Fig. 3C). in agreement with the correlations reported in various biological and experimental systems. including in Oc derived from the RAW264.7 cell line (An et al.. 2014; Ghazalpour et al.. 2011). In fact. among the 144 proteins of the Oc signature. only 72 genes were also in the Oc transcriptional signature. 2 were repressed (Table S3. pink and purple respectively) and 70 changed differentially or did not change significantly as compared to Dc and Mo. Similarly. among the 181 proteins of the immature Dc signature. 138 genes were also in the transcriptional signature and 5 were repressed (Table S4. pink and purple respectively). Not withstanding these discrepancies. RNAseq provides a much deeper view that SILAC of genome expression during Oc and Dc differentiation. For instance. no proteins were detected in SILAC for genes essential in Oc biological functions present in the Oc signature such as Siglec15 (Hiruma et al.. 2013) and Bcar1/p130Cas (Nagai et al.. 2013).

Finally. we examined the molecular function (MF) and biological process (BP) gene ontology (GO) terms associated with Oc and Dc signatures. All the proteins and all the genes identified by SILAC and RNAseq were respectively used as reference proteome and transcriptome. In the Oc protein signature. all enriched GO terms pointed at energy metabolism by the mitochondria (Fig. 4A and Table S7). in agreement with the up-regulation of tricarboxylic acid cycle enzymes during osteoclastogenesis and elevated energy metabolism in Oc (An et al.. 2014; Lemma et al.. 2016). Transcriptionally enriched GO terms in Oc also additionally comprised mRNA translation (Fig. 4B and Table S8). For Dc signatures. all enriched GO terms related immune functions (Fig. 4C-D and Tables S9-10). Consistent with the Oc GO term analysis. the main protein-protein interaction (PPI) networks among the Oc transcriptional signature were linked to mRNA translation and mitochondrial energy production (Fig. 4E-G).

**FIG. 4.**
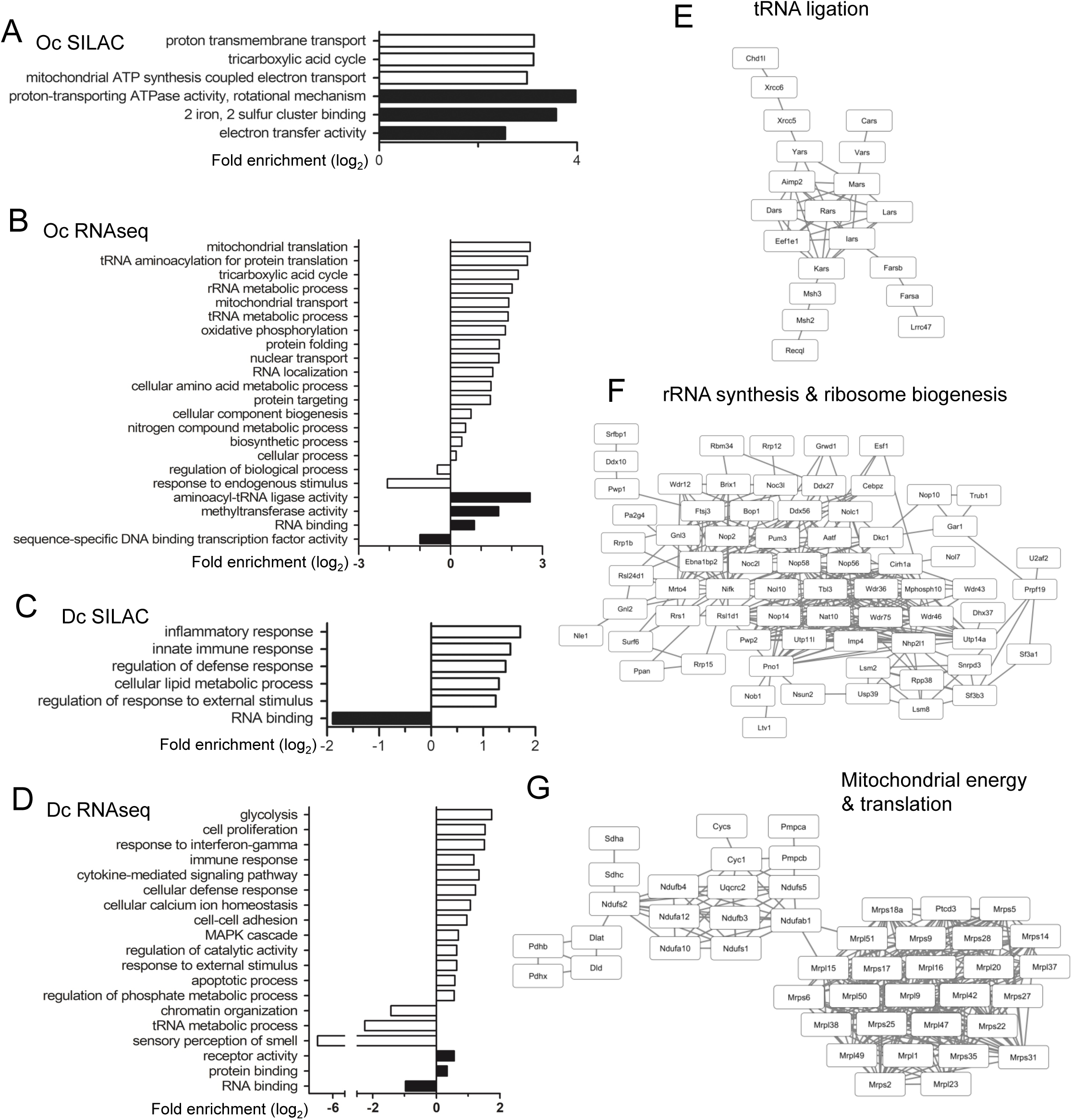
GO term enrichment analysis and protein networks. **(A-D)** Histograms represent log_2_ fold significant variations of GO terms linked to proteins (A-C) and genes (B-D) more abundant in Oc (A-B) and in Dc (C-D). White and bars correspond to GO terms associated with biological process and black bars to molecular function. GO term details are in Tables S7-10. **(E-G) Interaction networks among genes overexpressed in Oc**. Cytoscape representation of the amino acid to tRNA ligation (E). rRNA synthesis and ribosome biogenesis (E) and mitochondrial energy and translation (G) networks established with the RNASEQ data using STRING Protein-Protein Interaction Networks Functional Enrichment Analysis.

Overall. the proteomic and transcriptomic approaches in primary myeloid cells proved complementary to identify biologically relevant species characteristic of Oc and Dc. Moreover. the Oc and Dc signatures highlighted species of unknown function in these cell types. which could bring novel insights into the specific biological functions of these myeloid cells.

### Identification of potential new regulators of Oc biology

The 144 proteins of the Oc signature comprised 27 known regulators of Oc differentiation and activity (green highlight in Table S3) including cathepsin K and tartrate-resistant acid phosphatase type 5 (Cappariello et al.. 2014). the vATPase complex. a3 subunit Tcirg1 (Ochotny et al.. 2011) and Dock5 (Vives et al.. 2011; Vives et al.. 2015). other vATPase subunits. mitochondrial proteins and proteins involved mRNA translation. Interestingly. there were also 45 proteins of unknown function in Oc (light and dark blue highlight in Table S3). including proteins involved in cytoskeleton and intracellular trafficking. key processes for bone resorption (Cappariello et al.. 2014). Conversely. the 80 proteins less abundant in Oc comprised 7 known inhibitors of Oc differentiation or function (green highlight in Table S5) such as Pedf/Serpinf1. involved in human type VI osteogenesis imperfecta (Homan et al.. 2011). and Fkbp5 that was associated with human Paget’s disease (Lu et al.. 2017). So far. the other 73 proteins have not been linked to Oc biology and could comprise new Oc inhibitors (Table S5).

We then compared our data to the only SILAC global proteomic data available so far for Oc. concerning osteoclastogenesis in RAW 264.7 cells (An et al.. 2014). RAW 264.7 cell line was derived from the lymphoma of a mouse infected with Abelson murine leukemia virus; it can differentiate into Oc in the presence of RANKL. As shown by a recent SILAC study. the proteome of RAW 264.7-derived Oc is very distinct from that of primary Oc. in particular regarding cytoskeletal organization; unfortunately. as the data is not available. we could not compare it with ours (Ng et al.. 2018). 2.073 proteins were identified by SILAC during RAW 264.7 osteoclastogenesis (An et al.. 2014). among which 1.506 were common to our SILAC in primary Oc. They comprised 92 proteins of our primary Oc protein signature. but only 56 were up regulated during RAW264.7 Oc differentiation (Table S3. ↗). even though the log_2_ expression ratio cut off set at 0.5 in that study was less stringent than ours. The other proteins were down regulated or unchanged during RAW264.7 differentiation (orange highlight in Table S3. respectively ↘ and =). including known regulators of Oc biology induced during primary Oc differentiation such as Tensin 3 (Touaitahuata et al.. 2016) and CD68/macrosialin (Ashley et al.. 2011). Moreover. 52 proteins from our Oc signature were not detected in RAW 264.7 Oc (red highlight in Table S3. n.f.). Overall. our primary Oc signature comprised 89 proteins not highlighted in RAW264.7 Oc. SILAC proteome data available so far for Oc (An et al.. 2014). among which unknown regulators of osteoclast biology could be present.

These 89 proteins comprised: 15 known actors of Oc biology (green highlight in Table S3) and 36 proteins involved in biological processes activated in Oc (members of v-ATP complex. actors of mitochondrial energy metabolism and mRNA translation) and 38 proteins of unknown function in Oc (dark blue highlight Table S3). We picked about half of the remaining proteins for a siRNA screening. covering a variety of functions but avoiding the proteins with a role on chromatin (Tep1. Hmgb3. Histone H1.4 and H1.5) (bold lines in the dark blue proteins of Table S3). To avoid interference with the early differentiation process. primary Oc cultures were transfected at day 2 of differentiation with a commercial SmartPool of 4 siRNAs (Table S11) and grown for two more days to obtain Oc. We stained actin. tubulin and DNA. acquired mosaic images of entire wells on an automated microscope with a low-magnification objective and full-well images were reconstituted. The screen was performed in triplicate with 3 independent mouse Oc cultures. None of the 17 test siRNAs provoked cytoskeleton organization defects visible at that image resolution; moreover. no significant changes in Oc size were measured (not shown). Remarkably. with the 5 siRNAs targeting the AKR1C3 homolog Akr1c18. Dmxl1. Trip11. Vps52 and Wdr7 (Supp Fig. S5A). we observed a high proportion of Oc exhibiting very big and unusual intracellular structures delineated with actin and tubulin (Fig. 5A-B). The induction of Oc differentiation markers Src and cathespin K was not affected by the siRNAs (Fig. 5C-D). The “vacuole-like structures” had a regular circular shape and spanned over 10 *µ*m in all directions (Fig. 5E). We hypothesized that such structures may arise from the disturbance of Oc intracellular traffic. In fact. it was shown that the inhibition of lipid kinase PIKfyve. a regulator of the endolysosomal traffic induces the formation of large endolysosomes in RAW264.7 cells (Hazeki et al.. 2013). We did observe very large vesicles in Oc treated with PIKfyve inhibitor YM201636; these were positive for the endolysosomal marker CD63/Lamp3 and not delineated by actin (Supp Fig. S5B). This is in contrast with the vacuole-like structures elicited by the 5 siRNAs in Oc. which were surrounded by actin but were not delineated by any of the early and late endosomal marker we tested. including CD63 (Fig 5E). This suggests that these unusual actin and microtubule-delineated structures are not membrane compartments; we could not relate them to any subcellular compartment reported so far and they had no effect on the amount of cathespin K inside the Oc or secreted. nor on its maturation. as assessed by western blot (data not shown).

**FIG. 5.**
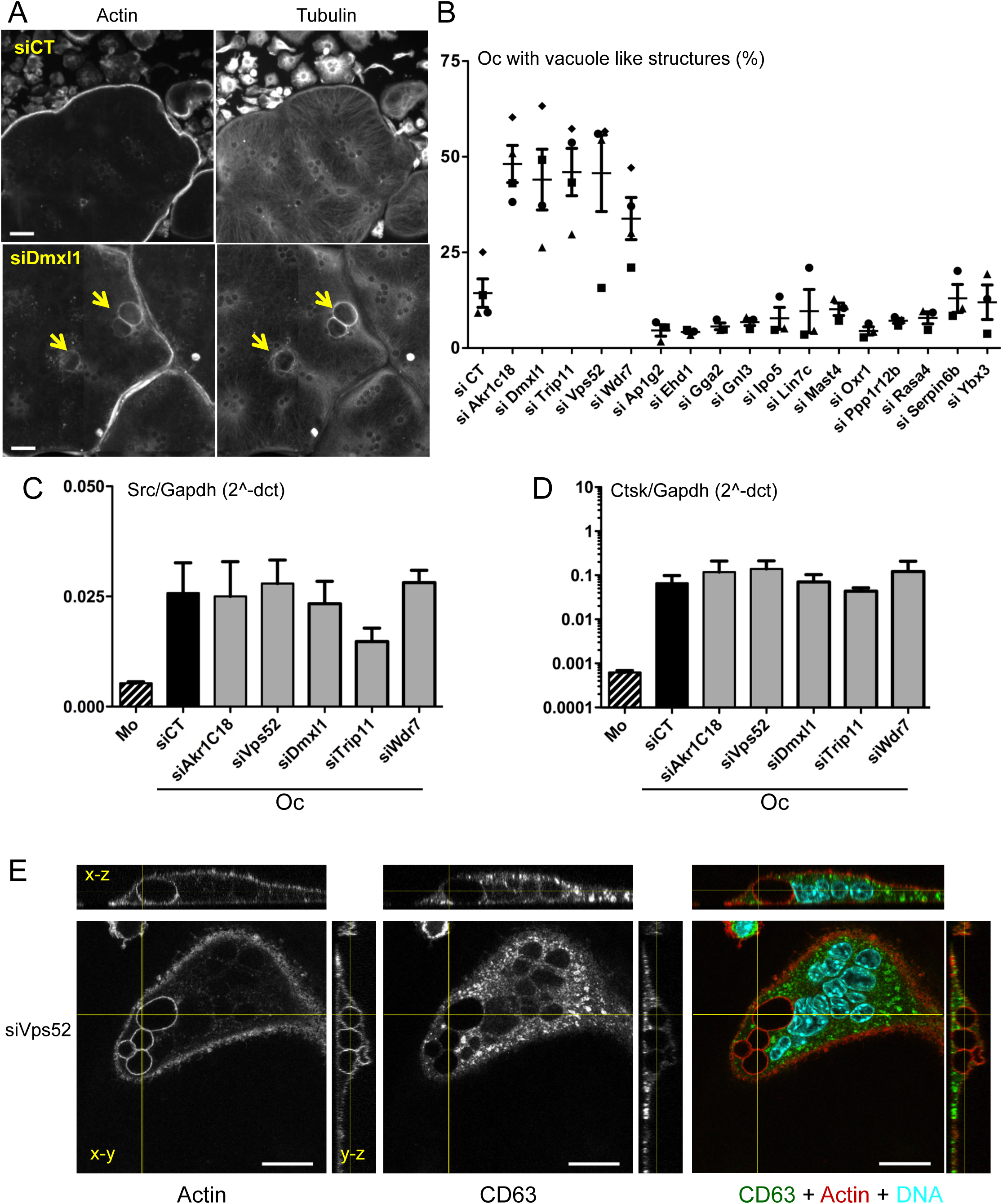
RNA interference of potential new regulators of Oc. **(A)** Examples of low-resolution images obtained by automated imaging and showing actin and tubulin staining in Oc transfected with control siRNA (siCT) and siRNA against Dmxl1. Arrows point at the large vacuole-like structures. Scale bar: 50 *µ*m. **(B)** Graph showing the proportion of Oc presenting large vacuole-like structures according to the siRNA transfected. counting at least 100 Oc per experiment and showing the results of 3 to 4 independent experiments depicted with a distinct symbol. with mean % and SEM. **(C-D)** Bar graphs show the mean and SEM mRNA levels Src (C) and cathepsin K (D) normalized to Gapdh. as measured by QPCR in bone marrow macrophages (Mo. dashed bars) and in Oc (black and grey bars) transfected with the indicated siRNAs. in 3 independent experiments. Efficiency of gene silencing is shown in Fig. S5A. **(D)** Confocal images representative of the large vacuole-like structures. from an Oc transfected with siRNA against Vps52. Panels show a single x-y confocal plan with the corresponding x-z and y-z views (46 plans with 377 nm z-steps) along the yellow lines. with staining for actin. CD63 and the overlay with DNA in the right panel. More plans Scale bar = 20 *µ*m.

These results suggest that our Oc signature comprises new regulators of Oc biology. which should be examined in more details to understand their functions.

### Tubb6 regulates Oc cytoskeleton and bone resorption

The SILAC and RNAseq data also revealed changes in the expression of ß-tubulins between Oc. Dc and Mo. Among all α and β tubulin genes. these cells express the same 4 α and 4 β isotypes: namely Tubba1a. 1b. 1c and 4a and Tubb2a. 4b. 5 and 6. Tubb5 (Tubulin beta class I) and Tubb6 (Tubulin beta 6 class V) represent more than 80% Tubb reads in the three cell types (Table S2). The protein levels of the 4 α and 4 β isotypes levels were not different between Oc. Dc and Mo. except for Tubb6 (Table S1). Indeed. Tubb6 showed 1.375 and 1.256 log_2_ protein folds as compared to primary Dc and Mo respectively. whereas such increase was not observed during RAW264.7 osteoclastogenesis (Table S3).

Tubb6 was reported to have a microtubule destabilizing effect in cycling cells. in contrast with Tubb5 (Bhattacharya and Cabral. 2009; Bhattacharya et al.. 2011). As microtubule dynamic instability is important for correct organization of actin and for bone resorption (Biosse Duplan et al.. 2014; Guimbal et al.. 2019). we explored the biological relevance of increased Tubb6 levels in Oc. Q-PCR analyses confirmed that among all ß-tubulin isotypes. Tubb6 expression uniquely increased during primary Oc differentiation (Fig. 6A). To specifically target Tubb6. we generated the 2 siRNAs si1077 and si1373 (Table S11). which neither affect the expression of Tubb2a. 4b and 5 RNAs nor the global Tubb protein level in Oc (Fig. 6B-C) and did not modify the expression of Oc differentiation markers Src and Ctsk (Fig. S6A-B). Interestingly. we observed that Tubb6 siRNAs strongly perturbed the organization of the podosome belt. provoking podosomes scattering at cell periphery (Fig. 6D-E). This was accompanied by a severe modification of microtubule morphology at cell periphery with the appearance of highly buckled microtubules (Fig. 6F). Using specific antibodies raised against the C-terminus of mouse Tubb6 (Spano and Frankfurter. 2010). we found that the distribution of Tubb6 was indistinguishable from that of pan-ß tubulin. except for a lower amount of Tubb6 in the mononucleated cells surrounding osteoclasts (Fig. 7G). consistent with increased expression of the Tubb6 during Oc differentiation (Fig. 6A).

**FIG. 6.**
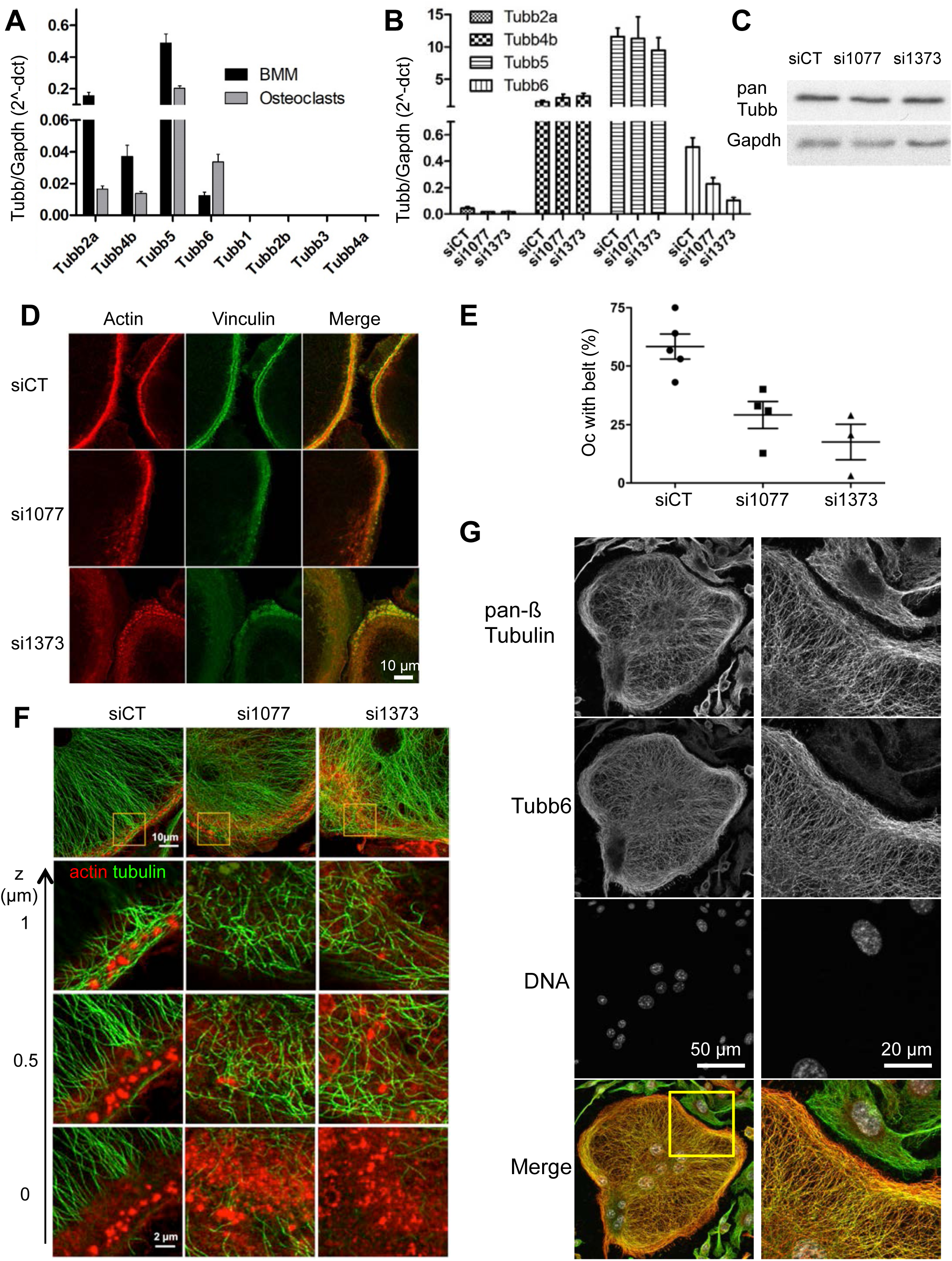
Role of ß tubulin isotype Tubb6 in Oc cytoskeleton organization. **(A)** Graph showing mean and SEM mRNA levels normalized to Gapdh of ß-tubulin isotypes in bone marrow macrophages (BMM. black bars) and Oc (grey bars). in three independent experiments and measured by QPCR. **(B)** Graph showing mean and SEM mRNA levels normalized to Gapdh of ß-tubulin isotypes expressed in Oc transfected with the indicated siRNA in five independent experiments and measured by QPCR. **(C)** Representative western blot showing total ß-tubulin and Gapdh expression in Oc transfected with the indicated siRNA. **(D)** Confocal images showing the detail of representative podosome belts in mouse Oc transfected with the indicated siRNA and labeled for actin (red) and vinculin (green). Scale bar: 10 *µ*m. **(E)** Bar graph shows the mean and SEM % of mouse Oc transfected with the indicated siRNA with a normal podosome belt. counting at least 100 Oc per experiment in 4 (si1077) and 3 (si1373) independent experiments. **(F)** Airyscan confocal images showing the detail of representative podosome belts in a mouse Oc transfected with the indicated siRNA and labeled for tubulin (green) and actin (red). Top panels show maximum projections of 13 plans with 172 nm z-steps. Zooms of the boxes areas in the top panels are shown below at different z. Scale bars: 10 *µ*m in top panels and 2 *µ*m in other panels. **(G)** Representative images of the localization of Tubb6 in a mouse Oc stained for pan-ß tubulin. Tubb6 and DNA. with boxed area enlarged in right column and image overlay in bottom row. showing no specific localization of Tubb6 (red) as compared to pan ß-tubulin (green) staining.

**FIG. 7.**
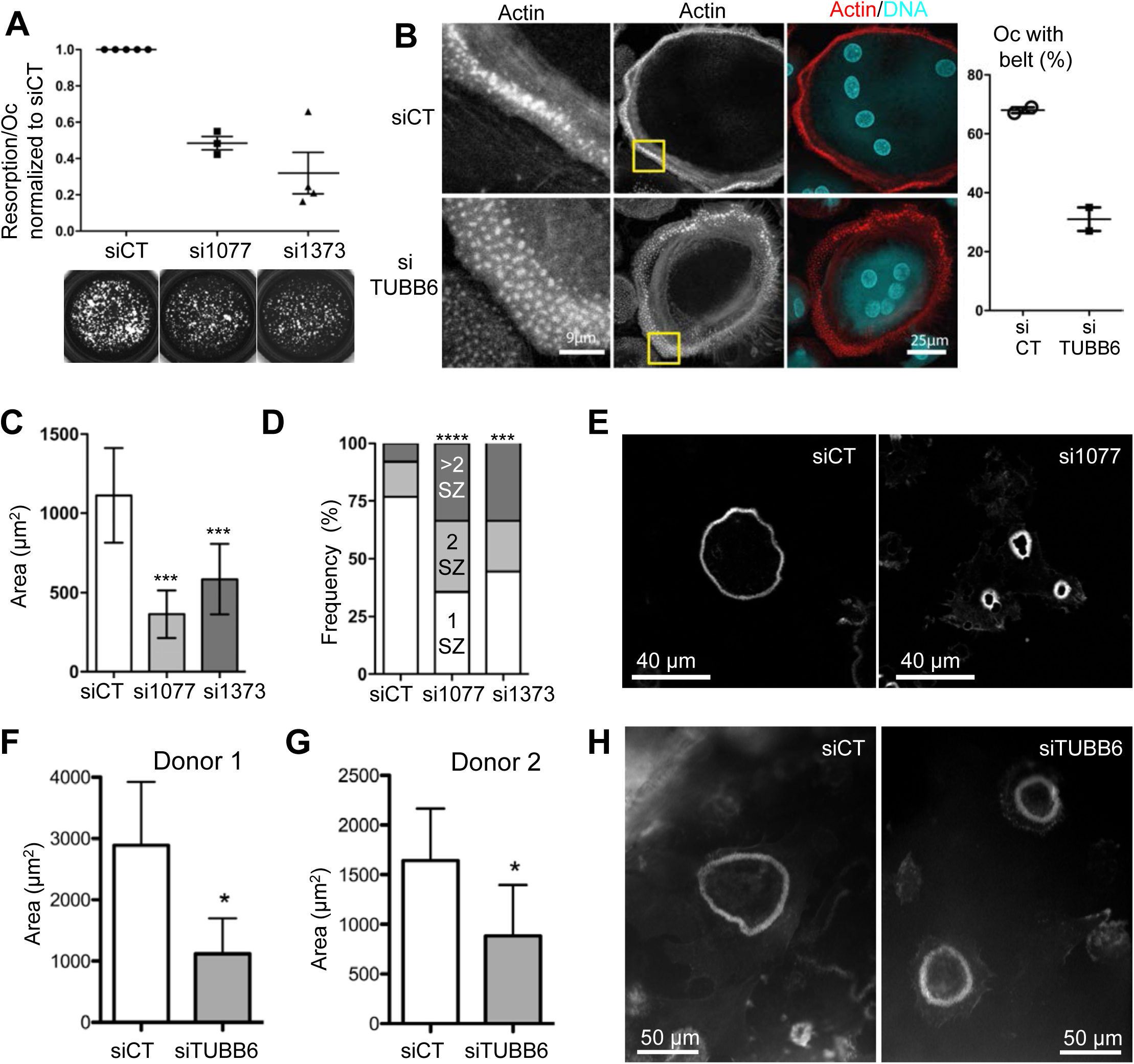
Effect of Tubb6 silencing on Oc activity and on the sealing zone of mouse and human Oc. **(A)** Bar graph comparing the mean and SEM specific resorption activity of Oc transfected with the indicated siRNA and normalized to control siCT. in 3 (si1077) and 4 (si1373) independent experiments. The von Kossa staining of representative wells of one experiment is shown below. **(B)** Confocal images (maximal projection of 7 plans. 210 nm z-step) showing representative podosome belts (middle column) and the details (left column) in a human Oc transfected with the control or TUBB6 siRNA and labeled for actin (red) and DNA (blue). Scale bar: 25 *µ*m. Bar graph shows the % of human Oc transfected with the indicated siRNA with a normal podosome belt. counting at least 100 Oc per experiment in 2 independent Oc preparations from CD14+ human PBMC. **(C)** Bar graphs shows the mean and 95% confidence interval of sealing zone area in mouse Oc sitting on ACC. measuring 96 (siCT). 133 (si1077) and 115 (si1373) sealing zones in one experiment ***: p<0.001. Kruskal-Wallis test with Dunn’s post test. **(D)** Bar graph show the distribution of the number of sealing zones (SZ) per mouse Oc in the same experiment. ***: p<0.001 and ****: p<0.001. Chi-square contingency. Result of a second experiment are shown in Fig. S6C-D. **(E)** Representative confocal images of the sealing zones measured in (A-B) in mouse Oc seeded on ACC and stained for actin. **(F-G)** Bar graphs shows the mean and 95% confidence interval of sealing zone area in human Oc sitting on bone slices in 2 independent Oc preparations from CD14+ human PBMC. measuring 14 to 19 sealing zones per condition; *: p<0.05. Wilcoxon two tailed test. **(H)** Representative confocal images of the sealing zones measured in (D-E) in human Oc seeded on bone and stained for actin.

Functionally. Tubb6 knock down resulted in severed reduction in the activity of Oc (Fig. 7A). Our former analysis of Affymetrix data of human myeloid cells also revealed increased expression of TUBB6 mRNA in Oc derived from human CD14^+^ peripheral blood monocytes (Gallois et al.. 2010; Maurin et al.. 2018). Thus. we examined the effect of TUBB6 siRNAs in human Oc differentiated form peripheral blood of 2 independent donors. Consistent with the results in mouse Oc. the proportion of human Oc presenting a normal podosome belt was also strongly reduced (Fig. 7B). We further analyzed the importance of Tubb6 on *bona fide* sealing zone. examining Oc plated on mineralized substrates. In mouse Oc plated on coverslips coated with collagen mineralized with calcium phosphate (ACC). we observed that Tubb6 siRNAs provoked the appearance of numerous and very small sealing zones per Oc instead of usually 1 or 2 bigger sealing zones in control Oc (Fig. 7C-E and Fig S6 C-D). This indicates that Tubb6 siRNAs induce a defect in sealing zone maturation. TUBB6 siRNAs also reduced the size of the sealing zones in human Oc differentiated form peripheral blood of 2 independent donors and plated on bone (Fig. 7F-H). These results suggest that higher levels of Tubb6 are important for efficient bone resorption. impacting on the organization of osteoclast podosomes and the structure of the sealing zone. Tubb6 overexpression was found to cause dystrophy in both human and mouse myotubes. correlated with expression changes of muscular heavy chain myosin genes (Randazzo et al.. 2019). Osteoclasts express two myosin heavy chain genes: *myh9* (myosin IIa) and *myh10* (myosin IIb) (Table S2). Myosin IIa. and not myosin IIb. is enriched at the podosome belt and the sealing zone; the silencing myosin IIa was found to increase sealing zone size (McMichael et al.. 2009). But in Oc treated with Tubb6 siRNAs. there was no modification of myosin IIa protein levels (data not shown).

Altogether. these results establish that the ß tubulin isotype repertoire is of functional importance in myeloid cells. In particular that the specific increase in Tubb6 protein levels that occurs during Oc differentiation is essential for microtubules to ensure correct patterning of Oc podosomes for efficient bone resorption.

## Discussion

We combined SILAC proteomics. RNAseq and homogenous experimental conditions to provide the first comparative global proteomes and transcriptomes of primary mouse Oc and Dc. From the data. we propose specific transcriptomic and proteomic signatures for each cell type. including proteins of unknown function that represent new candidate regulators of Oc and Dc biology. Using siRNA approach. we explored several of these new candidates in Oc. In particular. we found that Oc. Mo and Dc express the same repertoire of 4α and 4β tubulin isotypes. Within this repertoire. only Tubb6 protein is differentially expressed with higher levels in Oc. We further demonstrated that this increased expression of Tubb6 is essential for podosome organization and the control of sealing zone size in both mouse and human Oc. and that it is key for Oc resorption function.

The hematopoietic cells of the monocytic lineage are innate immune cells characterized by a great level of plasticity: mouse bone marrow macrophages and human CD14^+^ monocytes can differentiate in Dc and Oc; moreover. Dc have the capacity to transdifferentiate into Oc (Madel et al.. 2019). Still each cell type has very specific biological functions requiring particular cellular processes: Oc are the professional bone resorbing cells that participate in the maintenance of skeleton health whereas Dc are the sentinel antigen-presenting cells that sample their environment for “non-self” entities. Unraveling the molecular pathways underlying the biology of Oc and Dc is key to understand both their physiological and pathological roles. There are only few proteomic data sets available for Oc; furthermore. because of the difficulty to generate sufficient primary Oc for sample preparation. all were obtained using RAW264.7 cells as Oc precursors (Segeletz and Hoflack. 2016). RAW264.7 is a transformed macrophage-like cell line with osteoclastogenic potential. but this in an M-CSF independent manner; it was reported recently that there were major proteome discrepancies between primary Oc and Oc derived from RAW264.7 cells. in particular regarding cytoskeleton regulatory pathways (Ng et al.. 2018). Here we developed an experimental approach to prepare primary Mo. Oc and Dc amenable for quantitative SILAC proteomics. Thereby. we provide the first global quantitative proteomic dataset for primary Oc. In the present study. we focused our analyses on the Oc signature. Of note. the Dc signatures contain genes of unknown functions in these cells. including strongly differential genes that are poorly studied overall such as BC035044 or D630039A03Rik or Adgrg5. some of which could be relevant for Dc biology. For instance. Plet1 was recently shown to be involved in interstitial migration of murine small intestinal Dc (Karrich et al.. 2019).

To obtain the three myeloid cell types. we started from the same mouse bone marrow and used the same culture medium. just changing the cytokines added. The aim was to minimize the non-relevant changes that would not be related to the differentiation programs of the each cell types. To remain compatible with the technical requirements of our study. in particular the amount of material required for SILAC. we used plastic as a substrate for Oc. Dc and Mo. Although not their physiological environment. such procedure provides cells competent for their physiological functions. For instance. on plastic or glass. Oc have a ruffled border and they secrete active cathepsin K (Fuller et al.. 2010; Touaitahuata et al.. 2014b); when transferred onto bone. such Oc are able to resorb it. Still. previous reports showed that seeding Oc onto bone induces higher expression of various genes as compared to plastic (Crotti et al.. 2011; Purdue et al.. 2014). thus we may miss some of those genes in our Oc transcritpomic signature. These articles explicitly mention 25 genes whose expression levels are increased in Oc plated on bone as compared to plastic. Examining how these genes were classified in our study. we found that 12 genes were indeed in our Oc signature and 4 genes were not expressed (Table 1). Three genes were falling into our Dc signature and their fold induction on bone does not reflect genes characteristic resorbing Oc as compared to Dc. Finally for the remaining 6 genes. the induction fold was specified only for 3 genes: *Bikunin. Xdh* and *Max*. According to induction reported on bone and the expression we found in Oc. Mo and Dc. they may represent transcripts characteristic of the bone resorbing Oc absent from the Oc signature we established on plastic. *Xdh* and *Max* appear relevant in the bone resorbing Oc. Max (Myc-associated factor X) is a partner of Myc that modulates global gene expression. Myc was indeed shown recently to drives metabolic reprogramming during Oc differentiation and function. switching the metabolims to an oxidative state involving the TCA cycle and oxidative phosphorylation (Bae et al.. 2017). On the other hand. *Xhd* encodes the xanthine dehydrogenase (or xanthine oxidase) that participates in the generation of reactive oxygen species (ROS) through xanthine metabolism. The ROS generated by xanthine dehydrogenase have been shown to increase Oc formation and bone resorption (Fraser et al.. 1996; Garrett et al.. 1990). The gene *Bikunin. Ambp* in fact. is difficult to interpret as it encodes a precursor glycoprotein further cleaved into two proteins: the lipocalin alpha-1-microglobulin and the protease inhibitor bikunin (trypstatin). Overall. this confirms that Oc differentiation on plastic does recapitulate the transcriptional differentiation program of functional bone-resorbing Oc. Still. the transcriptional profile elicited by the transcription factor Max could be interesting to highlight genes specifically linked to the oxidative metabolism in the bone resorbing osteoclast.

**Table 1:** Analysis of the genes reported to upregulated when seeded on bone as compared to plastic.

| Gene | Bone/Plastic Fold (a) | Mean Oc FPKM (b) | Mean Dc | Mean Mo |
| --- | --- | --- | --- | --- |
| Oc signature |  |  |  |  |
| AnxA8 | 14.4 | 151.05 | 27.52 | 0.00 |
| Itgb3 | 7 | 6964.94 | 487.35 | 350.59 |
| CD82 | 9.4 | 4095.31 | 2757.66 | 2010.40 |
| Del1 (Edil3) | 2.6 | 2485.37 | 87.24 | 728.86 |
| Uchl1 | 1.69 | 4680.57 | 1320.44 | 1455.79 |
| Fkbp9 | 1.58 | 1903.45 | 241.65 | 1043.27 |
| Spns2 | 1.69 | 1181.08 | 175.55 | 11.66 |
| CD109 | n.s. | 2985.65 | 117.82 | 832.85 |
| Tm7sf1 (Gpr137b) | n.s. | 18291.29 | 3778.76 | 10596.39 |
| Mt3 | n.s. | 793.26 | 59.92 | 1.47 |
| CD44 | n.s. | 25674.43 | 17677.28 | 16530.46 |
| Rab38 | n.s. | 780.31 | 53.83 | 24.97 |
| Dc signature |  |  |  |  |
| Clec2 (Clec1b) | 5.5 | 14.73 | 233.22 | 14.12 |
| Timp3 | 4.4 | 1.00 | 24.75 | 2.20 |
| Sphk1 | 1.52 | 87.06 | 240.54 | 7.30 |
| Lower in Dc |  |  |  |  |
| Xdh | 4.8 | 6421.43 | 2555.57 | 8107.93 |
| Kif5b | n.s. | 6972.73 | 4337.10 | 7190.44 |
| Atp6v1a | n.s. | 25067.50 | 8107.01 | 19985.66 |
| Other |  |  |  |  |
| Odc1 | n.s. | 2226.47 | 2395.44 | 1120.10 |
| Bikunin (Ambp) | 3 | 13.53 | 3.83 | 34.96 |
| Max | 5.4 | 2305.00 | 2341.47 | 2165.10 |
| Not expressed |  |  |  |  |
| Prodh2 | 14.3 | 1.25 | 1.02 | 0.73 |
| Gabrq | 26.5 | 0.00 | 0.00 | 0.24 |
| Mpo | n.s. | 0.50 | 3.06 | 6.59 |
| AV253635 (C030013C21Rik lncRNA) | 7 | 0.25 | 0.26 | 0.98 |

(a) From Crotti et al., 2011 and Purdue et al., 2014 (Affymetrix Microarray data)
(b) Data from Table S2 including color code
n.s.: not specified, >1.5 after (Crotti et al., 2011)

We compared our primary Oc signature to the only SILAC global proteomics data reported for Oc. which was obtained from RAW264.7 cells (An et al.. 2014). Among the proteins of our primary Oc signature also detected in RAW264.7 Oc. a third were not changed or down-regulated during RAW264.7 osteoclastogenesis. including proteins induced during primary Oc differentiation and essential for bone resorption such as Tensin 3 (Touaitahuata et al.. 2016) and CD68/Microsialin (Ashley et al.. 2011). This confirmed the potential relevance of the proteins. but also likely the genes. in our primary Oc signatures that remain of unknown function. The siRNA screen revealed 5 siRNAs provoking the appearance of large vacuole-like structures in Oc. The target genes have diverse assigned functions: *Dmxl1* and *Wdr7* encode essential vATPase accessory proteins (Merkulova et al.. 2015). *Trip11* codes for GMAP-210 involved in vesicle tethering to Golgi (Roboti et al.. 2015). the product of *Vps52* is involved in endosome recycling (Schindler et al.. 2015) and *Akr1c18* is the mouse homolog of aldoketo-reductase *AKR1C3* involved in progesterone conversion (Lanišnik Rižner and Penning. 2019). Such intracellular structures delineated with actin and microtubules but without endosomal membrane marker were not described before; making it uneasy to hypothesize what molecular mechanism may be perturbed. We found that lipid kinase Pikfyve inhibition could induce very large intracellular structures of similar size in Oc but they were not delineated by actin and were endolysosomal membrane compartments. as reported in RAW264.7 cells (Hazeki et al.. 2013). More studies are needed on these potential new regulators of Oc biology. which would require stable knock down or knock out models to understand their precise functions.

The Oc signature also contained tubulin ß isotype Tubb6 (Tubulin beta 6 class V). In Oc. Mo and Dc. we found that the tubulin repertoire was restricted to the expression of only 4 α and 4 β tubulin genes; they do not express RNA for other tubulin isotypes such as Tubb1 that is needed for proplatelets elongation in megakaryocytes. another myeloid lineage hematopoietic cells (van Dijk et al.. 2018). This suggests the existence of a tight transcriptional control of the tubulin repertoire during hematopoiesis. but the mechanisms regulating of the transcription of tubulin genes remains very poorly understood (Gasic and Mitchison. 2019). How different tubulin isotypes affect microtubule dynamics to influence various processes in differentiated cells is an open question. The recent advances in producing recombinant tubulin isoforms started to shed light on the different biochemical properties of tubulin ß isotypes that can influence microtubule dynamics *in vitro* (Roll-Mecak. 2019). Tubb6 has the unusual property to destabilize microtubules in cycling cells whereas it is not the case for Tubb5 (Tubulin beta call I); this effect appears to rely specifically on the core domain of Tubb6 and not on its tail (Bhattacharya and Cabral. 2009; Bhattacharya et al.. 2011). Two aminoacids Ser239 and Ser365 in the core domain are critical for the microtubule destabilizing effect of Tubb6. and they are substituted for Cys in Tubb5. Interestingly. Tubb3 bears the Ser239 and Ser365 as in Tubb6; on the other had Tubb2b and Tubb5 have Cys residues at those positions. Biochemical studies revealed that Tubb3 and Tubb2b have distinct effects *in vitro*: on microtubule dynamics. on their association with microtubule associated proteins (MAP) as well as on their structure (Pamula et al.. 2016; Ti et al.. 2018). Therefore it is very likely that the level of Tubb6 can influence microtubule intrinsic properties as well as their dynamic behavior and association with MAP in Oc. Indeed. we found that the reduction of Tubb6 in Oc has a strong effect on microtubule and podosome organization. sealing zone size. resulting in impaired resorption activity. Contrarily to Tubb5 that is usually a major ß tubulin isotype with Tubb4b. the expression of Tubb6 is low in most tissues (Leandro-García et al.. 2010). and the tight control of Tubb6 expression levels appear key to various processes in differentiated cells. In fact. TUBB6 is among the most up-regulated genes in Duchenne muscular dystrophy (DMD) and persistent elevation of Tubb6 levels drives microtubule disorganization in the fibers of muscles in DMD mice (Randazzo et al.. 2019). TUBB6 levels also control the execution of pyroptosis by bacteria-infected lymphoblastoid cells (Salinas et al.. 2014). We show here that higher amount of Tubb6 in Oc is key for actin organization. sealing zone size and Oc resorption function. Recent reports highlighted the importance of microtubule dynamics and their cross talk with actin to control podosome patterning in Oc and bone resorption (Biosse Duplan et al.. 2014; Guimbal et al.. 2019; Zalli et al.. 2016). Thus. the enrichment of Tubb6 in Oc is very likely to participate in the molecular processes controlling actin and microtubule crosstalk to ensure efficient bone resorption. Whether it functions through affecting intrinsic microtubule dynamics. MAP binding or both remains to be elucidated.

In summary. we designed a homogeneous experimental set up to generate protein and RNA samples of primary Oc and Dc leading to relevant protein and transcript signatures of the two myeloid cell types. In particular. we found that high levels of ß tubulin isotype Tubb6 are characteristic of Oc and functionally relevant. Further studies on the specific properties of Tubb6 will bring better knowledge about the crosstalk between microtubules and actin cytoskeleton. which remains poorly understood. taking advantage of the unique functional context of bone resorption by Oc. This should be valuable also in the context of osteolytic bone diseases and pave the way for novel strategies against osteoporosis by targeting Oc cytoskeleton.

## Materials and methods

### Animals. human samples and ethics statement

4-week-old C57Bl/6J mice were purchased from Harlan France and maintained in the animal facilities of the CNRS in Montpellier. France. Procedures involving mice were performed in compliance with local animal welfare laws. guidelines and policies. according to the rules of the regional ethical committee. Monocytes from healthy subjects were provided by Etablissement Français du Sang (EFS). Toulouse. France. under contract 21/PLER/TOU/IPBS01/20130042.

### Production of bone marrow macrophages. osteoclasts and immature dendritic cells for SILAC and RNAseq

Bone marrow cells were extracted from long bones of C57BL/6J mice 6 to 8 weeks of age as described (Guimbal et al.. 2019) and cultured in α-minimal essential medium (αMEM. Lonza) containing 10% Fetal Bovine Serum (FBS. BioWest). heat inactivated at 56°C for 30 minutes. 2 mM glutamine (Lonza) and 100 U/mL penicillin and streptomycin (Lonza). 3 × 10^7^ cells per 150 mm dishes at 37°C. 5.5% CO_2_. in a humidified incubation for 24 hours. Non-adherent cells were then collected and used to differentiate the three myeloid cell types and prepare samples for stable isotope labeling by amino acids (SILAC) and RNAseq (Fig. 1A-B). For bone marrow macrophages (Mo) and osteoclasts (Oc). non-adherent cells were then plated 5 × 10^6^ cells per 10-cm plate in the same medium containing 100 ng/ml Macrophage colony-stimulating factor (M-CSF. Miltenyi) for 5 days. Cells were then detached using Accutase (SIGMA). washed in PBS and resuspended in Roswell Park Memorial Institute (RPMI) medium: SILAC RPMI 1640 (Lonza) supplemented with 10% dialyzed FBS (Life Technologies). 2.4 mM L-Prolin (SIGMA). 2 mM glutamine (Lonza) and 100 U/mL penicillin/streptomycin (Lonza). For Mo RNAseq and SILAC sample preparation. cells were then plated 0.8 × 10^5^ cell per well in 24 well plates in RPMI medium supplemented with 30 ng/ml M-CSF and with the L-enantiomers amino acids Lys (0.4 mM) and Arg (0.8 mM) (SIGMA) non-isotope-labeled (L amino acids). For Oc RNAseq sample preparation. cells were plated 1 × 10^5^ cell per well in 24 well plates in RPMI medium supplemented with 30 ng/ml M-CSF and 50 ng/ml Receptor Activator of NF-κB Ligand (RANKL. Miltenyi) and with L amino acids as above. For Oc SILAC samples. L amino acids were substituted for the same concentrations of heavy isotope–labeled (H) amino acids ^13^C_6_-^15^N_2_ L-Lys:2HCl and ^13^C_6_-^15^N_4_ L-Arg (Cambridge Isotope Laboratories). For immature dendritic cells (Dc) RNAseq sample preparation. cells were plated 4×10^5^ cells per well in 24 well plates in RPMI medium supplemented with 20 ng/ml Granulocyte Macrophage Colony Stimulating Factor (GM-CSF. Miltenyi) and with non-labeled amino acids as above. For Dc SILAC sample preparation. L amino acids were substituted for the same concentrations of intermediate isotope–labeled (M) amino acids 4.4.5.5-D4 L-Lys:2HCl and ^13^C_6_ L-Arg (Cambridge Isotope Laboratories). All media were changed every other day for 8 to 10 days. In these conditions. H and M amino acid incorporation in Oc and Dc respectively was over 95% and the Arg to Pro conversion rate was below 0.4%.

### Protein and RNA extraction

Following differentiation. Oc cultures were incubated with 120 *µ*l per well Accutase (SIGMA). for a few minutes to remove mono-nucleated cells. Then all cultures were rinsed once with PBS and processed for protein or RNA extraction. For SILAC samples. cells were lysed in 8M urea (SIGMA) and 10 mM HEPES (SIGMA). pH 8.0. 12 wells per cell type in a total of 400 *µ*l. Cell lysates were centrifuged at 20.000 × g for 2 minutes; protein concentration was determined using BCA reagent (ThermoFisher). protein concentration was around 25 *µ*g/mL per sample. Lysates were stored at −80°C until SILAC proteomic analysis. For each SILAC analysis. Oc. Dc and Mo protein samples derived from the same mouse. the experiment was done in triplicate with three independent mouse bone marrow extractions.

For RNAseq. total RNA was extracted and purified using RNeasy Mini Kit and QIAshredder spin columns (QIAGEN) according to manufacturer’s instructions. 8 wells for Dc. 12 well for Mo and 4 wells for Dc. to obtain around 10 *µ*g of RNA. RNA concentration was measured using a Nanodrop UV-visible spectrophotometer (Thermo Scientific) and RNA quality was assessed using a Bioanalyzer 2100 (Agilent Technologies). Lysates were stored at −80°C until RNAseq analysis. For each RNAseq analysis. Oc. Dc and Mo RNA samples derived from the same mouse. the experiment was done four times with four independent mouse bone marrow extractions.

### RNAseq analyses

The 12 RNA samples (2 *µ*g) were processed and analyzed in parallel by Fasteris SA (Switzerland). according to the “HiSeq Service Stranded Standard Protocol” (https://support.illumina.com/sequencing/sequencing_instruments/hiseq-3000.html). The stranded mRNA libraries were sequenced by HiSeq 4000 Illumina technology. generating single reads of 1x 50 bp. Adapter sequences were removed from the obtained 1x 50 bp reads and adapter trimmed reads were used for further analysis. About 30 million raw reads were obtained per sample (from 26.717.590 to 36.916.924). with around 99% of the reads mapping on reference mouse genome GRCm38. Multiple mapping percentages ranged between 23.62 and 32.43% according to sample (Fig. S1C). Sequence mapping (Mus musculus genome GRCm38. from iGenome downloaded on the 2017-07-13). normalization and estimation of transcript abundances (FKPM) were performed using the Tuxedo suite of short read mapping tools (Bowtie v2.0.5. Tophat v2.0.6. Samtools 1.2 and Cufflinks v2.1.1). Differential expression analysis was performed with DESeq2 R package from Bioconductor v2.13. For each comparison by pairs. the mean of the normalized counts obtained for the four replicates within each group of samples was calculated as well as the log_2_ fold change. The p. adjusted for multiple testing with the Benjamini-Hochberg procedure. was used to qualify fold changes as significant (padj<0.05).

### Mass spectrometry sample preparation. LC-MS-MS analysis and protein quantification

Within each of the 3 replicate SILAC experiments. equal amounts of Oc and Mo or Oc and Dc or Dc and Mo proteins were mixed. leading to a total of 9 samples that were processed in parallel. All MS grade reagent were from Thermo Fisher Scientific. grade Optima. For each sample. a total of 10 *µ*g of proteins were digested with 0.5 *µ*g LysC (Wako) in 600 *µ*l of 100 mM triethylammonium bicarbonate for 1 hour at 30°C. Then samples were diluted 3 times in the buffer and 1 *µ*g trypsin (Gold. Promega) was added overnight at 30°C. Peptides were then desalted using OMIX (Agilent) and analyzed online by nano-flow HPLC-nanoelectrospray ionization using a Qexactive HF mass spectrometer (Thermo Fisher Scientific) coupled to a nano-LC system (U3000-RSLC. Thermo Fisher Scientific). Desalting and preconcentration of samples were performed on-line on a Pepmap® precolumn (0.3 × 10 mm; Thermo Fisher Scientific). A gradient consisting of 0–40% B in A for 120 min (A: 0.1% formic acid. 2% acetonitrile in water. and B: 0.1% formic acid in 80% acetonitrile) at 300 nL/min. was used to elute peptides from the capillary reverse-phase column (0.075 × 500 mm. Pepmap®. Thermo Fisher Scientific). Data were acquired using the Xcalibur 4.0 software. A cycle of one full-scan mass spectrum (375–1.500m/z) at a resolution of 60.000 (at 200 m/z). followed by 12 data-dependent MS/MS spectra (at a resolution of 30.000. isolation window 1.2 m/z) was repeated continuously throughout the nanoLC separation.

Raw data analysis was performed using the MaxQuant software (version 1.5.5.1) with standard settings. Used database consist of mouse entries from UniProt (reference proteome UniProt 2016_11). 250 contaminants (MaxQuant contaminant database) and corresponding reverse entries. Each protein identified was associated to the corresponding gene in Mus musculus genome GRCm38. Relative proteins quantifications were calculated on the median SILAC ratios to determine log_2_ fold changes. In each comparison. proteins with |log_2_ abundance ratio| ≥ 0.75 were considered differentially abundant.

### Other bioinformatic analyses

Principal component analyses of both proteomic and transcriptomic results were made using R. software. For RNAseq. the hierarchical clustering analysis was performed in MeV tool v4.9 (Saeed et al.. 2003) using the four normalized expression counts for each cell type. Before analysis. values were normalized by rows using the software. An average linkage algorithm from a distance matrix calculated with the Pearson correlation coefficient was employed to make the analysis. Comparison between RNAseq and SILAC results and linear regressions were made using Graphpad Prism 5 software. GO term enrichment analyses were performed using Panther software (http://pantherdb.org/). The total lists of proteins and genes identified in this study were used as references. UniProtIDs and Ensembl IDs were employed as input type IDs for proteomic and transcriptomic results respectively. GO terms with p-value < 0.05 were considered significantly enriched. Histograms of enriched or diminished GO terms were represented using Graphpad Prism 5 software. Protein-Protein Interactions (PPIs) were determined using the Search Tool for Retrieval of Interacting Genes/Proteins (STRING) software v10.5 (https://string-db.org/) (Szklarczyk et al.. 2015). Only interactions based on experimental source with interaction score ≥ 0.7 were considered. PPIs networks were then represented using Cytoscape v3.6.1 (http://www.cytoscape.org/index.html) (Shannon et al.. 2003).

### Mouse Oc cultures for siRNA treatment and immunofluorescence

Mouse primary Oc were differentiated from bone marrow cells of 6-8-week-old C57BL/6J mice as described (Guimbal et al.. 2019). At day 2 of differentiation. siRNAs were transfected with siImporter in OptiMEM medium (Life Technologies) containing 30 ng/mL M-CSF and 50 ng/mL RANKL as described (Touaitahuata et al.. 2016) using 100 nM siRNA: either Dharmacon siGenome Smartpools or custom oligonucleotides from Eurogentec (Table S11). After 3 hours. medium was replaced with -MEM medium containing 30 ng/mL M-CSF and 50 ng/mL RANKL. When relevant. Oc were transferred after another 4 hours onto ACC-coated coverslips prepared as described (Maurin et al.. 2018). Medium was changed every other day during 3 to 4 days. until Oc differentiate. For Pikfyve inhibition. Oc were treated for 1 hour with 1 *µ*M YM201636 (Cayman) in 0.005 % DMSO.

Oc were then fixed in 3.2% paraformaldehyde and 10 *µ*M Taxol (SIGMA) in PHEM (Pipes 60 mM. Hepes 25 mM. EGTA 10 mM. MgSO4 4 mM. pH 6.9) for 20 min. For tubulin detection. cells were permeabilized for 1 min with 0.1% Triton X-100 in PBS and saturated with blocking buffer (1% BSA in PBS) before incubation for one hour with anti-α-Tubulin (T5168. SIGMA; 1:2.000) in PBS containing 1% BSA. For CD63 (LAMP-3). cells were permeabilized for 2 min with 0.1% Saponin in PBS and saturated with blocking buffer before incubation for one hour with antibody R5G2 (MBL Life Science; 1:200) in PBS containing 1% BSA. Alternatively. for Tubb6 localization. Oc were fixed for 10 minutes in Methanol at −20°C and saturated with blocking buffer (1% BSA in PBS) before incubation for one hour in PBS containing 1% BSA with anti pan-ß-Tubulin (E7. Developmental Studies Hybridoma Bank; 1:2.000) and anti-mouse Tubb6 antibody. a generous gift of Dr Frankfurter who generated and purified the antibody (Spano and Frankfurter. 2010). In all cases. cells were then incubated for one hour in Alexa-conjugated secondary antibodies and/or F-actin marker Alexa-conjugated Phalloidin (Life Technologies; 1:1.000) or Rhodamin-labeled Phalloidin (SIGMA; 1:10.000) in PBS containing 1% BSA and bisBenzimide Hoechst dye (SIGMA) to stain DNA when relevant.

For siRNA screening. images were acquired at low resolution directly in the 24-well plates with an automated ArrayscanVTi microscope (Thermo) equipped with a 10X EC Plan Neofluar 0.3NA objective as described (Maurin et al.. 2018). For confocal imaging. samples were mounted in Citifluor mouting medium (Biovalley) and images were acquired with an SP5 confocal microscope equipped with Leica LAS-AF software with objectives 40X HCX Plan Apo CS 1.3 NA oil or a 63X HCX Plan Apo CS 1.4 NA oil (Leica). or a Zeiss Axioimager Z2 microscope equipped with MetaMorph 7.6.6 Software (Molecular Devices) with objectives 40X EC Plan Neofluar 1.3 NA oil or 63X Plan Apochromat 1.4 NA oil (Zeiss). For Airyscan confocal. imaging was performed with a 63x Plan Apo 1.4NA of a Zeiss LSM880 confocal microscope equipped with an Airyscan detector (32 GaAsp detector) in super resolution mode; Alexa 488 was excited at 488nm with argon laser and Alexa 647 at 633nm with helium/neon laser; Airyscan analysis was made using Zen software with default settings.

### Human Oc cultures for siRNA treatment and immunofluorescence

For differentiation to human Oc. monocytes purified by CD14^+^ purification kit (Miltenyi) were seeded on slides in 24-well plates at a density of 5.10^5^ cells per well in RPMI supplemented with 10% FBS (SIGMA). M-CSF (50 ng/mL) and RANKL (30 ng/mL); the medium was then replaced every 3 days with medium containing M-CSF (25 ng/mL) and RANKL (100 ng/mL) (Miltenyi) until Oc differentiation as described (Raynaud-Messina et al.. 2018). Targets silencing was performed at day 5 using reverse transfection protocol as previously described (Troegeler et al.. 2014). Shortly. cells were transfected with 200 nM of ON-TARGETplus SMARTpool siRNA targeting TUBB6 (Table S11) or the ON-TARGETplus Non-targeting control pool (Dharmacon) using HiPerfect transfection system (Qiagen) in RPMI. Four hours post-transfection. cells were incubated for 24 hours in RPMI-1640 medium. 10% FBS. 20 ng/ml of M-CSF and RANKL (100 ng/mL) for 7 additional days. Alternatively. Oc were transferred onto bone slices after 4 days and cultured another 3 days. Then. cells were fixed with PFA 3.7%. Sucrose 30 mM in PBS After permeabilization with Triton X-100 0.3%. saturated with blocking buffer. washed and incubated with Alexa Fluor 555 Phalloidin (33 mM. Thermo Fisher Scientific) and DAPI (500 ng/mL. SIGMA) in blocking buffer for 1 hour. Coverslips were mounted with Fluorescence Mounting Medium (Dako). Imaging was performed with a SP5 confocal microscope as above. For human Oc sealing zones. bone slices were imaged with a Leica DM-RB fluorescence microscope or on a FV1000 confocal microscope (Olympus) as described (Raynaud-Messina et al.. 2018).

### Image quantification. Oc activity assays. Q-PCR and western blot

The podosome belt status of Oc was determined considering that the podosome belt was abnormal when. in over half of Oc periphery. actin staining was fragmented and/or weak and/or thin and/or absent and/or that podosomes were scattered as described before (Maurin et al.. 2018). The measurement of sealing zone size were performed with ImageJ 1.51w software. Total RNA and protein extractions. real time Q-PCR and western blots were performed as reported previously (Maurin et al.. 2018) using the primers described in Table S11. primary antibodies against pan ß tubulin (sc-398937 1:1000. Santa Cruz) and Gapdh (#2118 1:10.000. Cell Signalling) and horseradish peroxidase-conjugated anti-mouse or anti-rabbit secondary antibodies (respectively NA931V and NA934V. 1:10.000. GE Healthcare). Mineral dissolution activity of Oc was measured as described (Maurin et al.. 2018). Briefly. at day 3 of differentiation (24 h after siRNA transfection). Oc were detached with Accutase (SIGMA) for 5–10 min at 37 °C and seeded for 3 days onto inorganic crystalline calcium phosphate (CaP)-coated multiwells (Osteo Assay Surface. Corning). 8 wells per siRNA. For each siRNA. four wells were then stained for Tartrate Resistant Acid Phosphatase (TRAP) activity to count Oc and four wells stained with Von Kossa to measure CaP dissolution as described (Brazier et al.. 2009) and imaged with a Nikon SMZ1000 stereomicroscope equipped with a Nikon DXM 1200F CCD camera. Quantification of resorbed areas per wells was done with ImageJ 1.51w software. In each experiment. Oc specific activity was expressed as the average area resorbed in the 4 wells stained with von Kossa normalized by the average number of the Oc in the 4 wells stained with TRAP. Statistical analyses were performed with GraphPad Prism 5.0. All imaging was performed at the Montpellier Ressources Imagerie (MRI) imaging facility (www.mri.cnrs.fr) except human Oc sealing zone images. which were made at the TRI-Genotoul imaging facility (trigenotoul.com).

## Acknowledgements

We acknowledge the imaging facility MRI. member of the national infrastructure France-BioImaging supported by the French National Research Agency (ANR-10-INBS-04. “Investments for the future”). We whish to thank Dr Anthony Frankfurter. Virginia University. Charlottesville. USA. for his generous sharing of Tubb6 antibodies. We are very grateful to Frank Comunale. Christelle Dantec. Philippe Fort and Peggy Raynaud. CRBM Montpellier. France. for helpful advice and discussions during transcriptomic analysis and for reagent sharing.

## Main figure Legends

**Figure S1.**
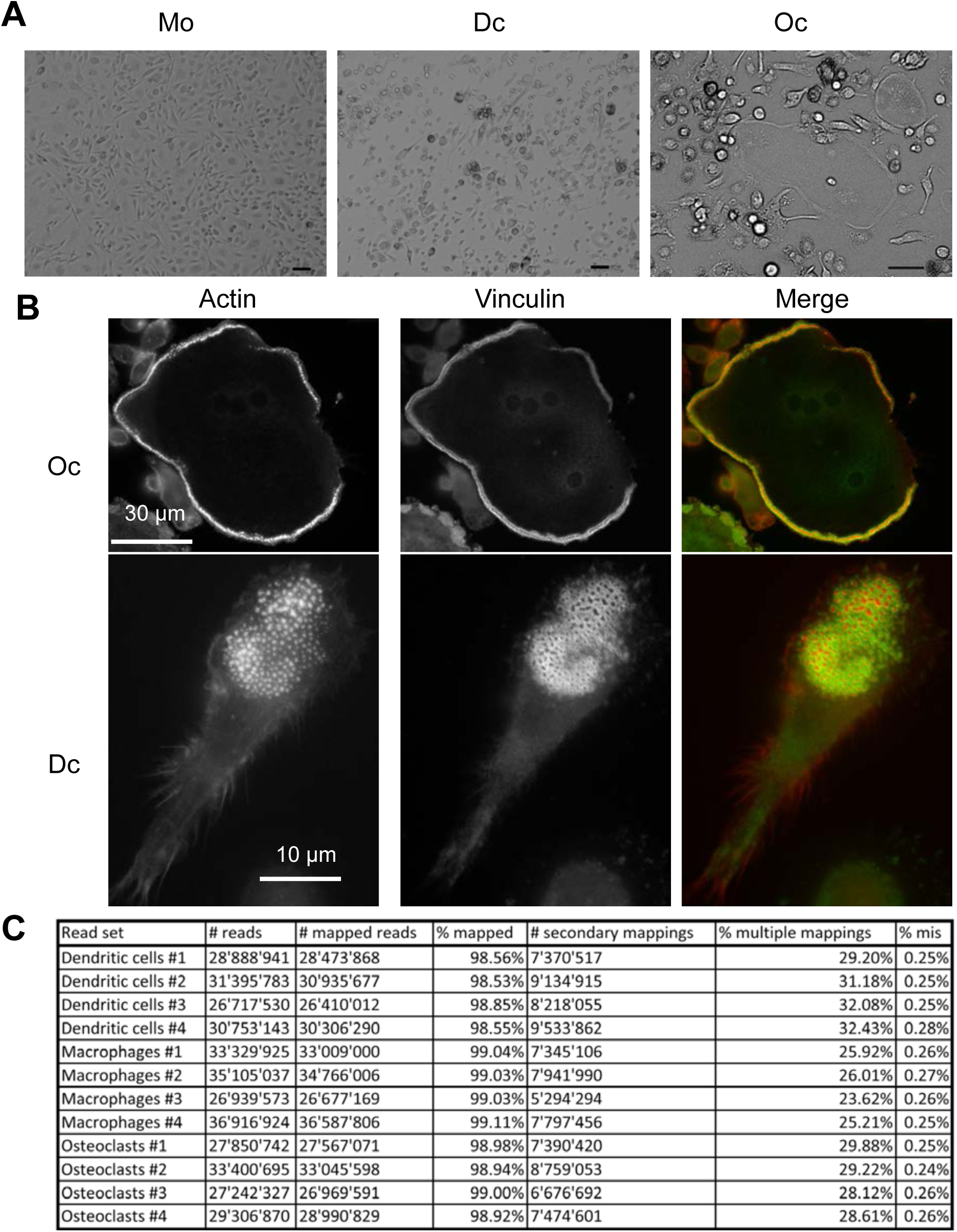
Differentiation of myeloid cells from mouse bone marrow cells in RPMI medium. **(A)** Phase-contrast images of Oc, Dc and Mo differentiated from bone marrow cells in RPMI medium. Scale bars: 50 *µ*m. **(B)** Wide field fluorescence images of Oc and Dc labeled for actin and vinculin. **(C)** RNAseq mapping results for each RNA sample.

**Figure S2.**
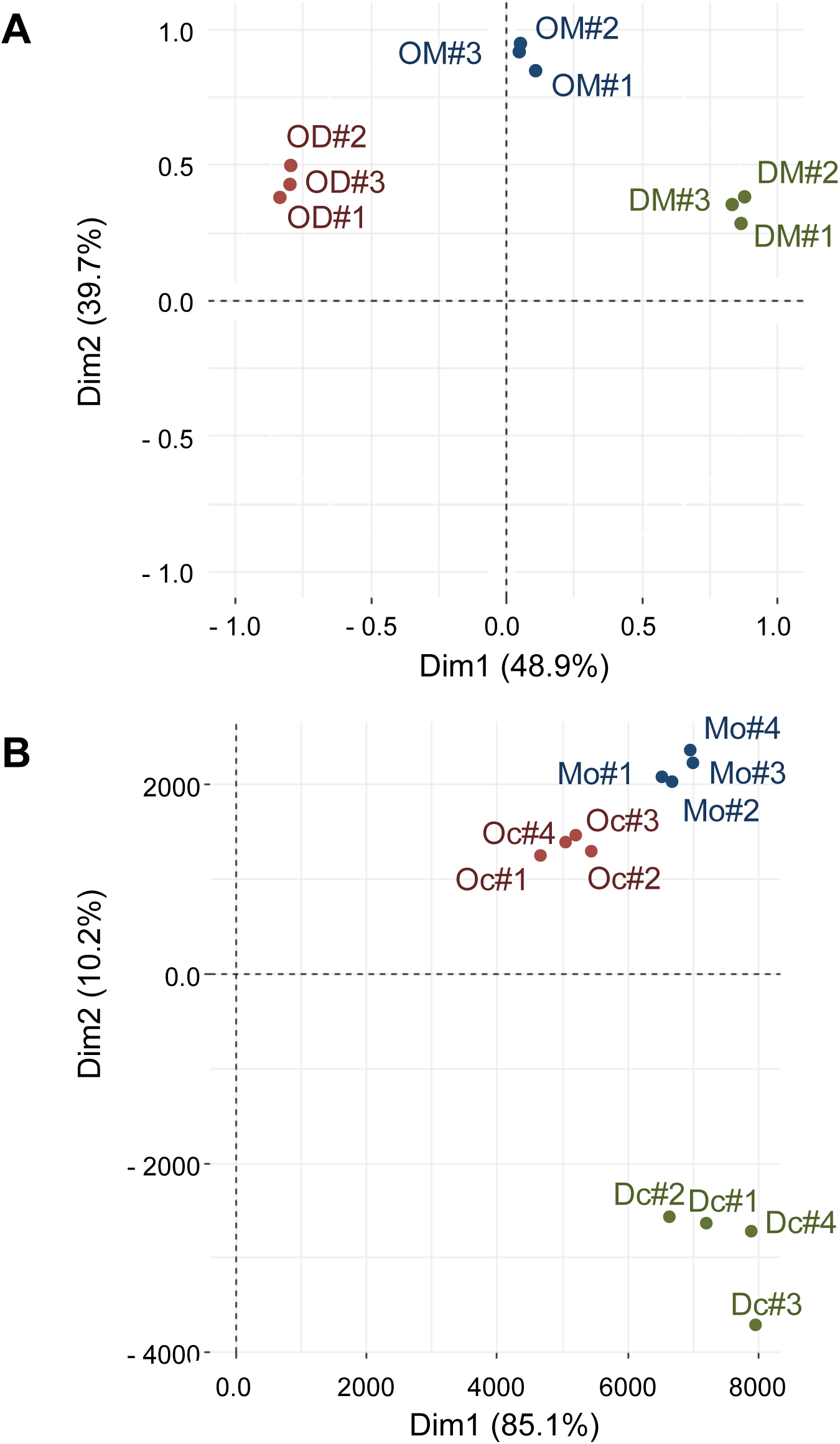
Sample validation and outlier detection in proteomic and transcriptomic studies. **(A-B)** Graphical representation of the 2-dimension Principal Component Analysis (PCA) of **(A)** the three samples in the three SILAC experiments and **(B)** the three samples in the four RNAseq experiments. In (A): OM: Oc vs Mo, DM: Dc vs Mo and OD: Oc vs Dc.

**Figure S3.**
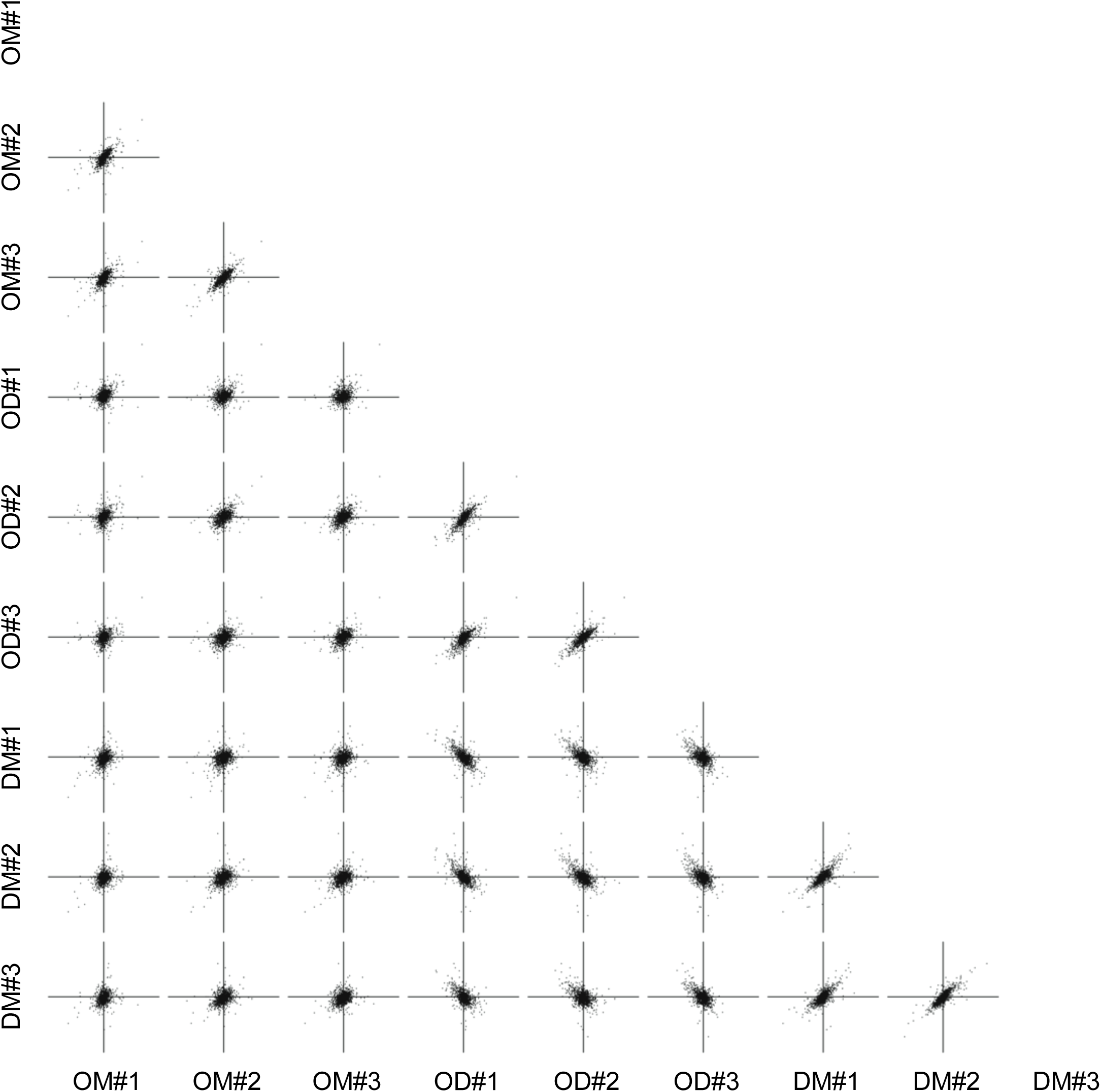
Correlation between protein samples. Graphs show the comparison of protein log2 abundance ratios between the three samples in the three SILAC experiments. OM: Oc vs Mo, DM: Dc vs Mo and OD: Oc vs Dc.

**Figure S4.**
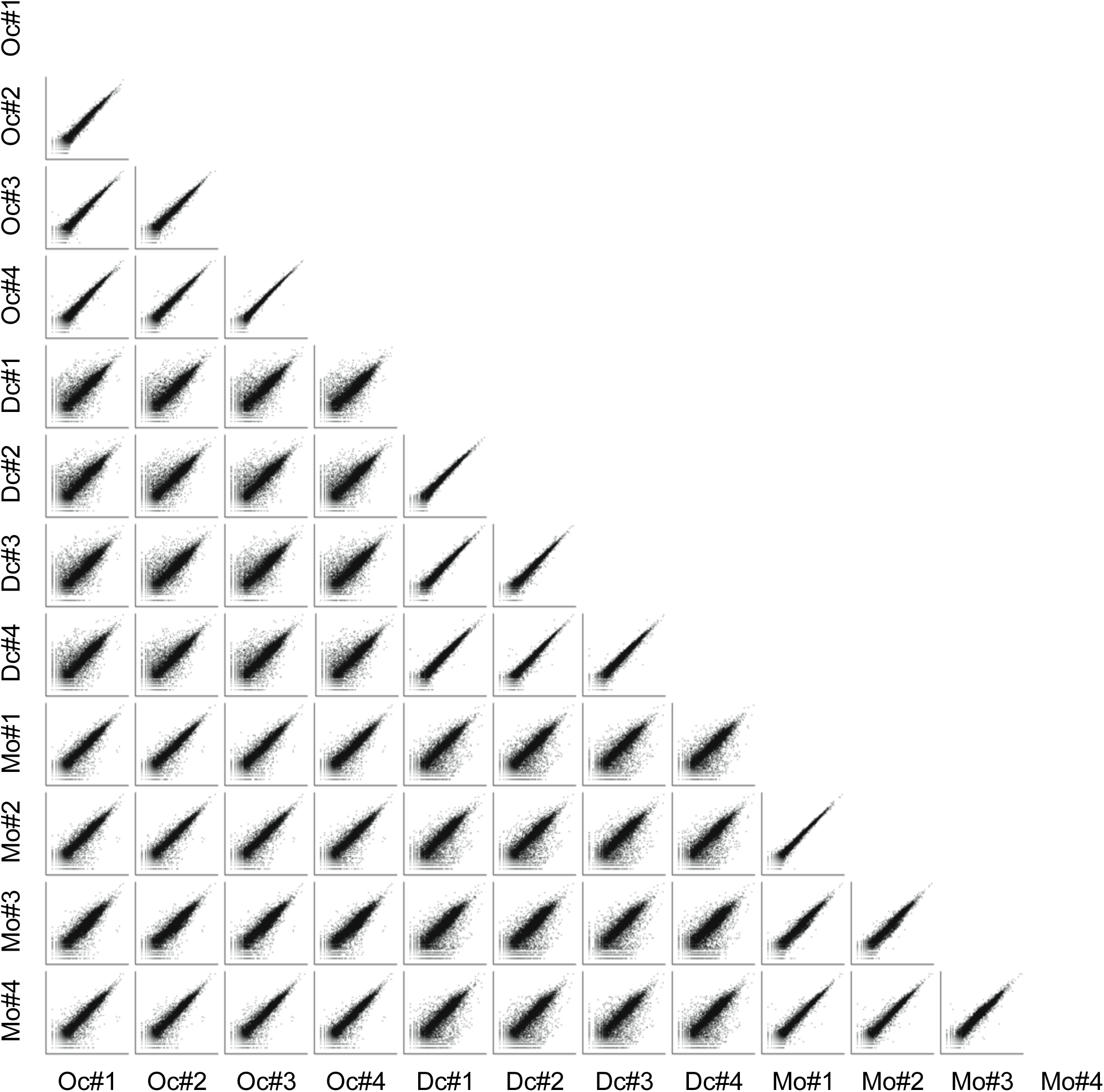
Correlation between RNA samples. Graphs show comparison of gene log_10_ FPKM between the three samples in the four RNAseq experiments.

**Figure S5.**
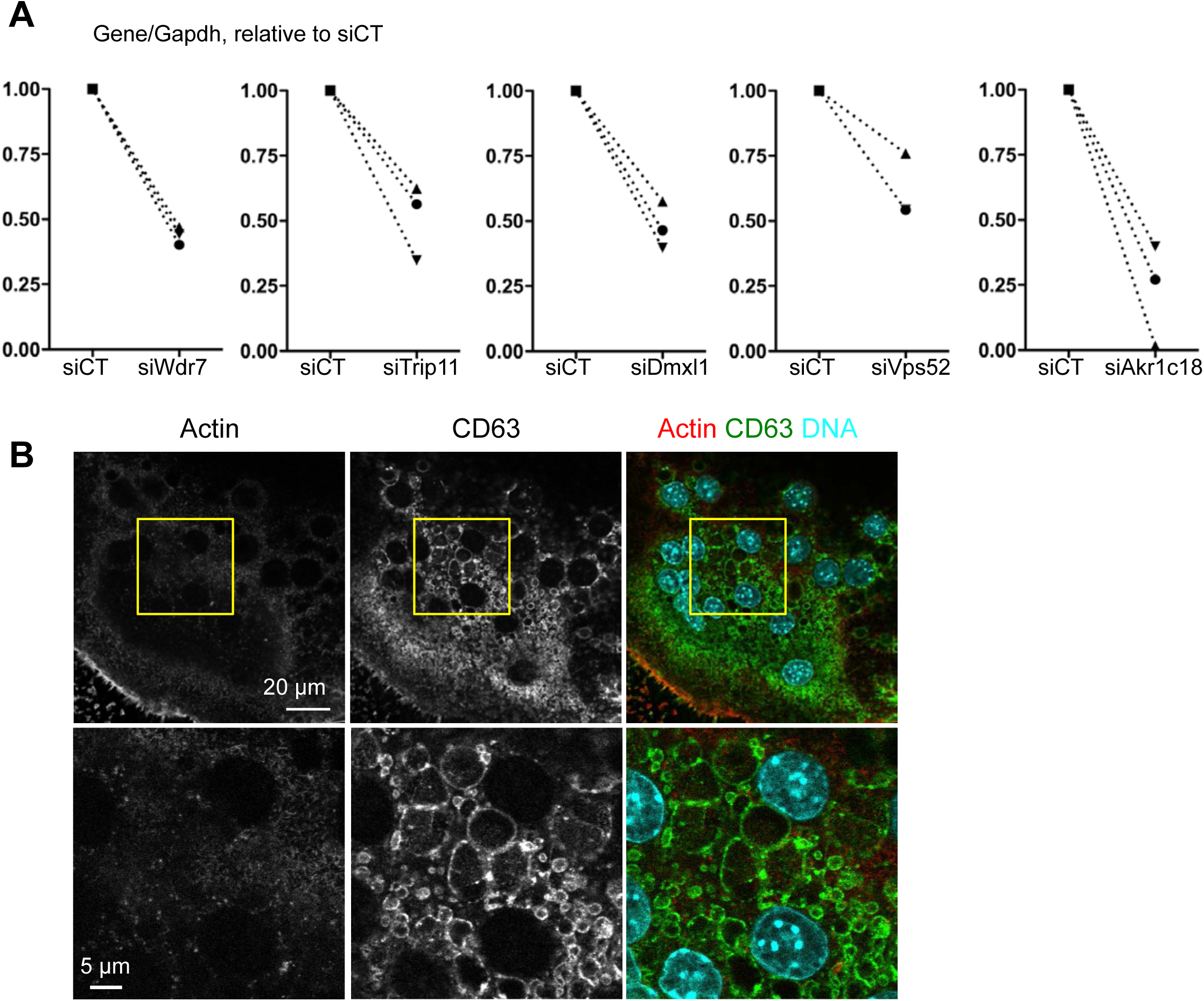
**(A) Efficiency of gene silencing.** The expression of Wdr7, Trip11, Dmxl1, Vps52 and Akr1C18 was determined by RT-PCR using the primers described in Table S11. The measurements were performed in three independent Oc differentiations from three mouse bone marrow preparations, each symbol representing one experiment. In each experiment, data were normalized to gene expression levels in Oc treated with the control siRNA. **(B) Effect of Pikfyve inhibition in Oc**. Representative confocal image of an Oc treated with 1 µM YM201636 for 1 hour showing a single x-y confocal plan (top panels) and magnification of the boxed area (bottom panels) with staining for actin, CD63 and the overlay with DNA. Scale bars: 20 µm (top panels) and 5 µm (bottom panels).

**Figure S6.**
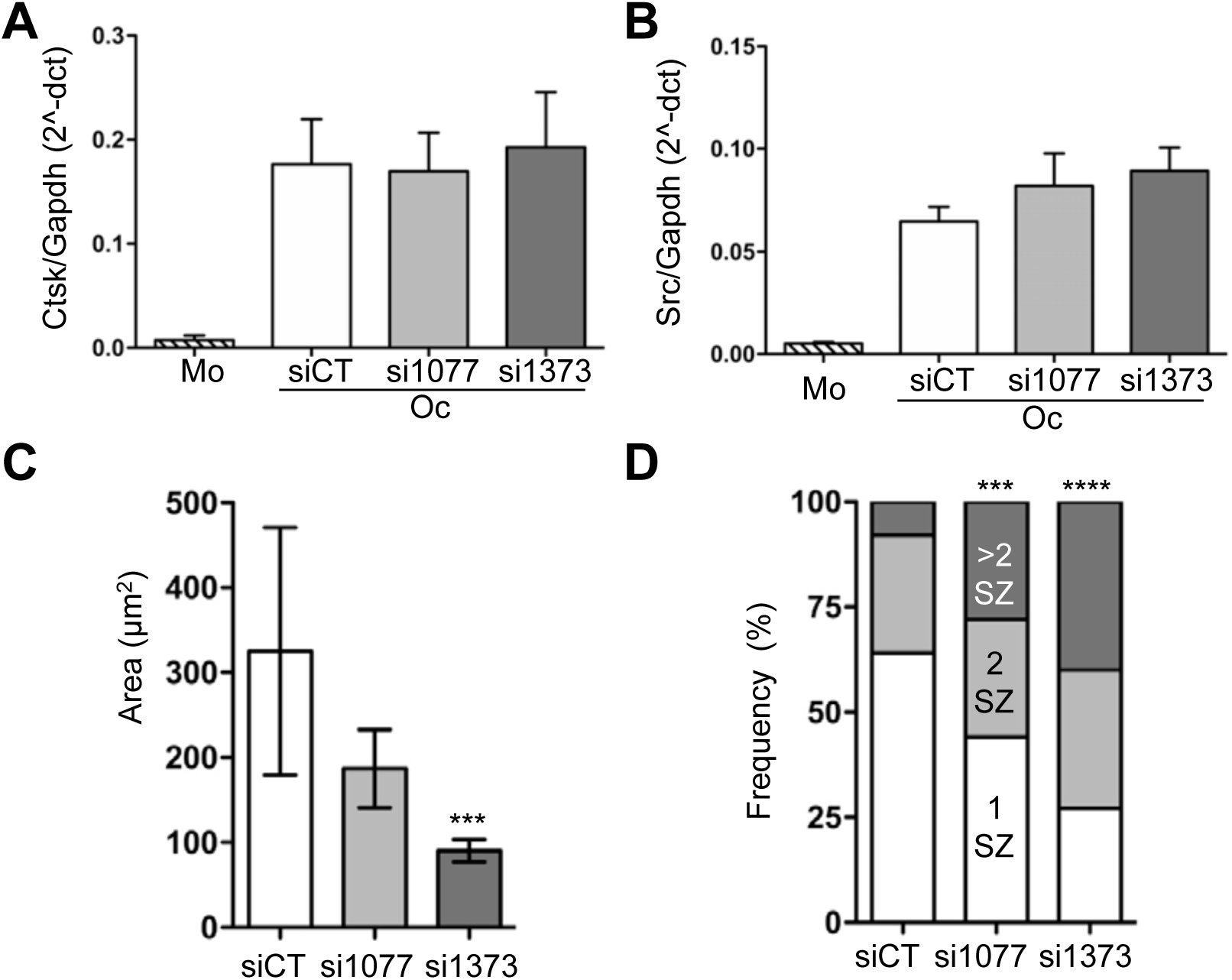
**(A-B) Q-PCR on Oc markers.** Bar graphs show the mean and SEM mRNA levels normalized to Gapdh as measured by QPCR of cathepsin K **(A)** and Src **(B)** in bone marrow macrophages (Mo) and in Oc (black and grey bars) transfected with the indicated control (black) and Tubb6 (grey) siRNAs, in 3 independent experiments. **(C)** Bar graphs shows the mean and 95% confidence interval of sealing zone area in mouse Oc sitting on ACC, measuring 60 (siCT), 58 (si1077) and 110 (si1373) sealing zones in one experiment; ***: p<0.001, Kruskal-Wallis test with Dunn’s post test. (B) Bar graph show the distribution of the number of sealing zones (SZ) per mouse Oc in the same experiment, ***: p<0.001 and ****: p<0.001, Chi-square contingency.

Table S1: List of the 2820 non-redundant proteins identified using SILAC proteomics

Table S2: List of the 17,199 genes identified using RNAseq

Table S3: Proteins more abundant in osteoclasts

Table S4: Proteins more abundant in dendritic cells

Table S5: Proteins less abundant in osteoclasts

Table S6: Proteins less abundant in dendritic cells

Table S7: Enriched GO terms linked to proteins more expressed in osteoclasts

Table S8: Enriched and diminished GO terms linked to genes more expressed in osteoclasts

Table S9: Enriched and diminished GO terms linked to proteins more expressed in dendritic

Table S10: Enriched and diminished GO terms linked to genes

Table S11: siRNAs and oligoniucleotides used in this study

